# Neuronal extracellular vesicles mediate BDNF-dependent dendritogenesis and synapse maturation via microRNAs

**DOI:** 10.1101/2021.05.11.443606

**Authors:** Anna Antoniou, Loic Auderset, Lalit Kaurani, Andre Fischer, Anja Schneider

**Affiliations:** Institute for Neurodegenerative Diseases and Geriatric Psychiatry, University Hospital Bonn, Germany; German Center for Neurodegenerative Diseases (DZNE), Bonn, Germany; Department of Psychiatry and Psychotherapy, University Medical Center Göttingen, Göttingen, Germany; Department for Systems Medicine and Epigenetics in Neurodegenerative Diseases, German Center for Neurodegenerative Diseases (DZNE), Göttingen, Germany

## Abstract

Extracellular vesicles (EVs) have emerged as novel regulators of several biological processes, in part via the transfer of EV content such as microRNA; small non-coding RNAs that regulate protein production, between cells. However, how neuronal EVs contribute to trans-neuronal signaling is largely elusive. We examined the role of neuron-derived EVs in neuronal morphogenesis downstream signaling induced by brain-derived neurotrophic factor (BDNF). We found that EVs perpetuated BDNF induction of dendrite complexity and synapse maturation in naïve hippocampal neurons, which was dependent on the activity of three microRNAs, miR-132-5p, miR-218 and miR-690. These microRNAs were up-regulated in BDNF-stimulated EVs. Moreover, supplementation with BDNF-EVs rescued the block of BDNF-induced phenotypes upon inhibition of miRNA activity. Our data therefore suggest a major role for EVs in BDNF-dependent morphogenesis, and provide new evidence for the functional transfer of microRNAs between neurons. This is not only an important step towards understanding the function of EVs in inter-neuronal signaling, but is also relevant for many disorders characterized by decreased BDNF signaling, such as major depression or cognitive impairment.

## Introduction

Extracellular vesicles (EVs) are lipid membrane-enclosed vesicles that have recently emerged as important regulators of development, adaptation and homeostasis in several biological systems^1^. In the brain, EVs were shown to mediate communication between neurons and non-neuronal cells such as oligodendrocytes^2,3^, astrocytes^4,5^, microglia^6–8^ and vascular endothelial cells^9^, and have been implicated in the spreading of misfolded, aggregating proteins such as tau^10–12^, amyloid-beta^13,14^ and alpha-synuclein^15,16^ in neurodegenerative disease models. An increasing number of reports demonstrate a function of EVs in neurobiological processes, namely neurogenesis^17^, excitatory synapse pruning^18^ and inhibitory neurotransmission^19^. Nevertheless, the precise contribution of EVs in neuronal morphogenesis is still largely elusive.

Although small EVs were previously commonly referred to as exosomes, which originate from the fusion of multi-vesicular endosomes (MVE) with the plasma membrane, it is now established that most EV preparations are heterogeneous^20,21^. For simplicity, we therefore classify EVs based on their size rather than biogenesis. Small EVs (sEVs; ~50-200nm) from many different cell types were shown to contain non-coding RNAs (ncRNAs), such as microRNAs (miRNAs)^20,22^, which are potent inhibitors of cytoplasmic protein production and regulators of gene expression^23–25^. Neuronal miRNAs play key roles in neuronal morphogenesis and are crucial regulators of synaptic plasticity; that is experience dependent changes in the strength of synapses^25^. Interestingly, evidence of functional miRNA transfer between cells has previously been reported in brain cells. For instance, neuron-secreted miR-124 upregulates the glutamate transporter GLT1 in astrocytes^5^, and neuronal EV-miR-132 was shown to promote vascular integrity by regulating vascular endothelial cadherin in neuroepithelial cells^9^. Moreover, it was previously reported that EV-miRNAs secreted at the synaptodendritic compartment are predicted to target regulators of neurite outgrowth^26^, and miRNAs and other small ncRNAs were shown to be present in synaptic vesicles^27^. These and other publications in other biological systems ^28–30^ indicate that miRNA regulatory networks may extend beyond cellular barriers. Whether neurons communicate via EV-miRNAs is currently unexplored.

We hypothesized that EVs play a role in neuronal morphogenesis within the context of BDNF signaling. BDNF has important roles in neuronal development, plasticity and neuromodulation^31^, in part by regulating local translation at neurites^32^, and miRNA production and activity^33–35^. In turn, several miRNAs were shown to mediate BDNF-dependent processes, for instance by targeting the actin regulator Limk1^36^ and the major regulators of transcription and translation, CREB^37^ and Pumilio^38^, respectively. Our data show that BDNF selectively regulates the sorting of specific miRNAs in sEVs. EVs from BDNF- but not control-stimulated neurons increased dendrite morphogenesis and induced the clustering of the pre-synaptic marker synaptophysin at dendrites of naïve hippocampal neurons. We further showed that these effects were dependent on the synergistic activity of BDNF-regulated EV-miRNAs. Importantly, defects in BDNF-dependent morphogenesis upon inhibition of these miRNAs was rescued by EVs from BDNF- but not control-stimulated neurons. Overall, this data demonstrates a specific role for neuronal EVs in BDNF-dependent dendrite maturation, and is to our knowledge the first report of functional inter-neuronal miRNA transfer.

## Materials and Methods

### Plasmids

Plasmids for dsRed, membrane GFP (mGFP) and dual fluorescence miRNA sensors were used as previously described (Antoniou et al 2018). Custom primers were purchased from Sigma and miRNA probes were from Life Technologies (*Supplementary Methods*, *Table S1*).

### Cell Culture

Unless otherwise stated cell culture media solutions were purchased from Invitrogen. Cell lines were not used after 25 passages and regularly tested for mycoplasma contamination by PCR.

#### Primary neuronal cell culture

Primary cortical and hippocampal neurons were derived from NMRI mice at embryonic day 16 (E16) according to Animal Welfare regulations. Pregnant mice were obtained from Charles River Laboratories (Sulzfeld, Germany). Brain tissue was dissected in HBSS medium supplemented with 1mM sodium pyruvate, 0.1% glucose and 10mM HEPES (pH7.3) and neurons were dissociated using 0,025% Trypsin. Neurons were seeded onto cell culture dishes (Falcon) or nitric-acid washed glass coverslips (Carl Roth GmbH) coated with 0,5mg/ml poly-D-lysine (Sigma) in plating medium (Basal medium Eagle’s containing Eagle’s salts, 0.45% glucose, 10% horse serum (Capricorn; HOS-1A), 1mM sodium pyruvate, 100U/ml Penicillin and 0,1mg/ml Streptomycin. Plating medium was replaced with serum-free maintenance medium MEM supplemented with 0.6% glucose, 0,2% sodium bicarbonate, 1mM sodium pyruvate, 2% B27 supplement (Gibco), 2mM Glutamax, 100U/ml Penicillin and 0,1mg/ml Streptomycin. Neurons were kept in a humidified incubator supplied with 5% CO_2_ at 37°C.

#### Cell Culture Treatments

Recombinant BDNF (Peprotech, #450-02) was dissolved in sterile water containing 0.1% bovine serum albumin (BSA). Neurons were treated with 50ng/ml BDNF or control vehicle for 20-30 minutes unless otherwise stated, and washed out 3 times with maintenance medium. Recombinant myc and myc-BDNF were from Chromotek and Cusabio, respectively. For blocking MAPK activity, neurons were treated with 10μM U0126 (Merck Millipore) for 30 min before addition of BDNF or control vehicle. Dynasore (Sigma-Aldrich) was dissolved in DMSO and used at 40μM, 30 min prior to addition of EVs. GW4869 (Sigma) was used at 5μM. AraC (Sigma) was used at the final concentration of 2μM on 1 DIV and washed out 24 hours later. Lipofectamine 2000 reagent was used to transfect primary hippocampal neurons and N2A neuroblastoma cells according to manufacturer’s instructions. Transfections were performed at 5 DIV, 1-2 days prior to BDNF or EV treatments.

### EV isolation and treatments

For all experiments, fresh, serum-free collection medium was added to donor cells 16 hours before EV isolation. Unless otherwise stated, 5-7DIV primary cortical neurons were used as EV donors and 6-7DIV hippocampal neurons were EV recipients. For functional experiments, sEV stock concentration was 150ng EV protein /μl (EVs from 100,000 donor cells /μl). Unless specified, a final concentration of 3μg EV protein /ml was used.

For more information on EV isolation, fluorescent labeling and NTA see *Supplementary Methods*.

### RNA Isolation

Total RNA was isolated using peqGOLD Trifast (VWR) or miRNeasy micro kit (Qiagen). Genomic DNA contamination was eliminated using either TURBO DNA-free kit (AM1907) or on-column digestion with RNAse-free DNAse set (Qiagen), respectively. RNA concentration was quantified using Nanodrop spectrophotometer.

### Real-time quantitative PCR (rt-qPCR)

Reverse transcription was carried out using iScript cDNA synthesis kit (170-8891, Bio-Rad) for mRNA targets and TaqMan Advanced miRNA cDNA synthesis kit (ThermoFischer; A28007) for miRNAs. PCRs were performed on StepOnePlus Real-Time PCR system (Applied Biosystems) using either iTaq SYBR Green Supermix with ROX (Biorad, #172-5121) or TaqMan Fast Advanced Master Mix (ThermoFischer, #4444557) for TaqMan assays. For NGS validation, miScript II RT kit synthesis was used for cDNA synthesis and candidate miRNA-specific primers were used in SYBR Green-based PCR.

### Small RNA sequencing

Twelve small RNA libraries representing 2 biological and 3 technical replicates from control- or BDNF-treated cortical neurons and corresponding EVs were prepared using NEBNext_Multiplex Small RNA Library Prep Set for Illumina (New England Biolabs) as per manufacturer’s instructions and sequenced in Illumina HiSeq2000. An in-house developed pipeline was used to analyze the small RNAome. Quality check and demultiplexing were performed using the CASAVA 1.8.2 software (Illumina). To quantify small RNAome, reads were first mapped to mature miRNA sequences obtained by miRBase (http://www.mirbase.org/) followed by further mapping to other small non coding RNA sequences (http://www.ensembl.org/info/data/ftp/index.html). Reads were then mapped to the mm10 reference genome. Target prediction and gene set enrichment analysis was preformed using miRWalk (https://www.mirwalk.umm.uni-heidelberg.de/).

### Immunofluorescence

Neurons were fixed in 4% paraformaldehyde/ 4% sucrose/ 1x phosphate buffered saline (PBS) solution for 15-20min at room temperature. For immunocytochemistry experiments, cells were permeabilized in 0.1% Triton-X100 for 5 min, washed in PBS and immunostained with primary and secondary antibodies diluted in 2% Bovine Serum Albumin (BSA) / 0.1% Tween-20 / 1xPBS. To image EVs in recipient cells, permeabilization was performed using 0.25% Saponin/ 5% BSA/ PBS for 30 min at room temperature, and the same solution was used for primary and secondary antibody incubation. Primary antibodies were added onto glass coverslips for either 1 hour at room temperature or overnight at 4°C, and secondary antibodies were incubated for 45-60 min at room temperature in a light-protected, humidified chamber. Coverslips were mounted on imaging slides using Mowiol 4-88 solution containing DABCO (24mg/ml) and DAPI (1:10,000). Images were acquired on Zeiss LSM880 confocal microscope (DZNE LMF).

### Quantification of dendrite complexity

XY scans of dsRed-expressing neurons were acquired using a 20x objective and dendrite complexity was quantified using Sholl analysis. Concentric circles were placed around the neuronal soma at 15μm intervals (end radius of 190μm), and the number of intersections between the circles and neuronal dendrites was counted in thresholded images using the Sholl analysis plugin in Image J. The number of intersections was quantified for approximately 10 neurons per condition and averaged for each independent experiment.

### Dual-fluorescence sensor assay

N2a cells were co-transfected with sensor plasmids and 10nM LNAs and EVs were added 24 hours later. Cells were fixed after approximately 22-24 hours. XY images were acquired using a 20x objective with picture tiling on two regions of interest (ROI). Sensor repression was quantified by randomly selecting 50-60 GFP-expressing cells and calculating the proportion of cells co-expressing dsRed. GFP cells were selected blindly, prior to red channel visualization.

### Synapse quantification

XY images of dsRed-expressing neurons were acquired at optimal resolution settings using a 63x objective. For quantification of pre- and post-synaptic marker levels, mean intensity was calculated at 20×8μm ROIs placed at random on primary and secondary dendrites of similar thickness. Colocalization analysis was performed using thresholded ROIs in the Coloc2 plugin in Image J. For puncta analysis, PSD95- and SYP-positive puncta were counted in dendritic spines using dsRed as a morphological marker. Only clearly defined puncta at the tip of dendritic spines were included in analysis. For statistical analysis, the average of 10 neurons and 2-3 dendrites per neuron was calculated for each experimental trial.

### Quantification and data analysis

Data are represented as mean ± standard deviation, unless otherwise stated. Statistical significance tests were performed as indicated in the figure legends using GraphPad Prism software. All imaging experiments were performed in a blinded manner.

## Results

### BDNF does not affect EV yield

We first characterized EVs secreted from primary mouse embryonic cortical neurons at 5-6 days in vitro (DIV). To exclude EVs secreted at earlier time points, conditioned maintenance media was washed out and replaced with fresh culture media 16 hours prior to EV isolation. Cell culture supernatants were then subjected to differential centrifugation to clear debris (3,500×*g*) and large vesicles (4,500×*g*) and to isolate medium-sized EVs (mEV; 10,000 ×*g*) and sEVs (100,000 ×*g*) (**Fig. 1A**). Using nanoparticle tracking analysis (NTA) we quantified the size distribution of re-suspended sEV and mEV pellets, which peaked at 168.5 ± 5.6nm and 196.6 ± 8.7nm in size, respectively, with mean particle concentrations of 3.66e8 ± 9e7 and 2.17e8 ± 6.22e7 particles/ml, respectively (**Fig. 1B**). The size distribution of particles derived from cleared 3,500 ×*g* supernatants peaked at 159.1 ± 4.1nm (**Fig. 1C**), suggesting that the majority of extracellular particles are indeed small in size. Importantly, non-conditioned cell culture medium processed in parallel had negligible particle counts in NTA, confirming that the medium itself does not account for isolated EVs (**Fig. 1C**). Moreover, inhibition of exosome secretion by the neutral sphingomyelinase (N-SMase) inhibitor GW4869 decreased particle concentration by approximately 1.5-fold in both sEVs and cleared supernatants and did not affect the mEV fraction (**Fig. S1A-B**). We further validated the size of sEV preparations using scanning transmission electron microscopy (STEM), which had a mean size of 150.5 ± 13.5nm under control conditions (**Fig. 1D**, **S1D**). Furthermore, we confirmed the purity of EV fractions in western blot (**Fig. 1E**). Here, we observed the presence of luminal endosomal proteins Alix, TSG101 and Flotilin-2 and the transmembrane protein Lamp1 in sEV fractions, which did not contain the Golgi marker, Grp75, or the endoplasmic reticulum protein Calnexin. We did not observe strong signals of EV proteins in mEV fractions, likely due to the lower yield of mEVs as shown by NTA measurements (**Fig. 1B**). Even though embryonic neuronal cultures grown in the absence of serum should not contain a significant amount of glia cells, contamination with glia-derived sEVs could lead to misinterpretation of our data. We therefore examined whether depletion of mitotic cells from neuronal cultures using cytosine d-D-arabinofuranoside (AraC) affects sEV yield. To validate glia cell depletion, we immunostained primary cortical neurons with antibodies against the neuronal marker MAP2 and the astrocyte-specific marker GFAP, which revealed that the vast majority of cells are in fact neuronal (**Fig. S1H**). Moreover, treatment of neuronal cultures with AraC, depleted GFAP signals and had no effect on the yield or size distribution of sEVs quantified by NTA (**Fig. 1G**). As AraC affects the long-term viability of neuronal cultures, which may itself influence the neuronal secretome, we did not use it in subsequent experiments.

**Figure 1:**
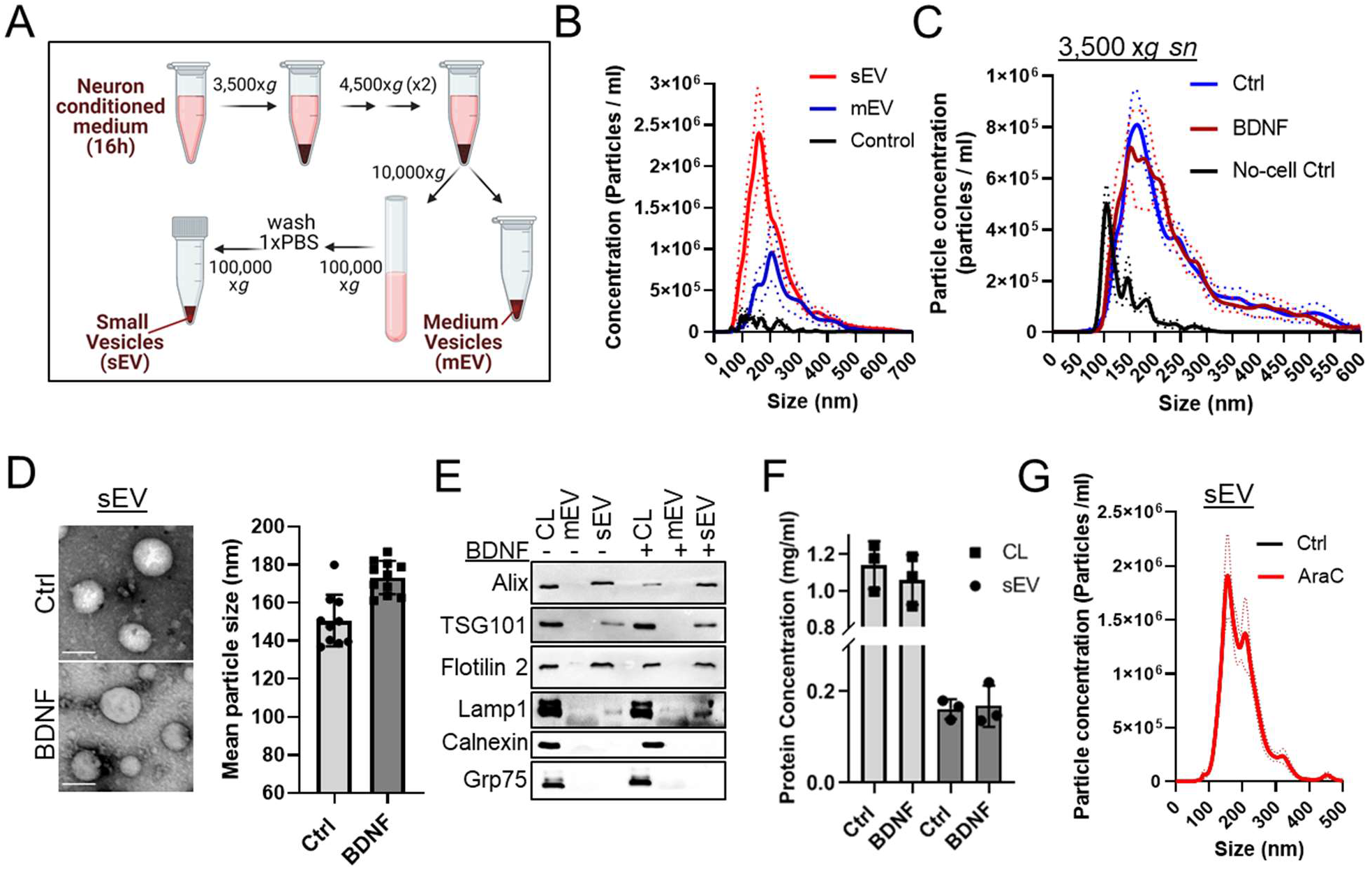
BDNF does not affect EV secretion. **A)** Diagram depicting isolation protocol for small and medium-sized EVs (sEV, mEV). Figure was created in biorender.com. **B)** Size distribution of sEVs and mEVs derived from primary cortical neurons, measured in nanoparticle tracking analysis (NTA). The buffer used to dilute the EV pellets was used as control; n=3-4, error bars represent standard error of the mean (dashed line). **C)** BDNF treatment does not affect the yield of extracellular particles. Cleared cell culture supernatants (3,500 ×g sn) from ctrl or BDNF-treated cortical neurons (7DIV) and non-conditioned culture medium (‘no-cell ctrl’) were processed in NTA. Traces and dotted lines represent the mean and standard error from four independent experiments, n=4. **D)** Scanning transmission electron microscopy (STEM) images of sEVs obtained from Ctrl or BDNF-treated neurons; scale bars are 100nm. (*Right*) Mean particle size of sEVs was calculated in approximately 10 STEM micrographs for each condition (dimensions 1,14 μm^2^). **E)** BDNF does not change the relative abundance of EV markers. Cell lysates (CL) and respective sEV and mEVs were processed in western blotting. The endoplasmic reticulum protein calnexin and golgi marker grp75 were used as negative controls. **F)** BDNF does not change total protein concentration in cortical neuron lysates (CL) or corresponding sEVs isolated by UC; n=3. **G)** Depletion of glia in neuronal cultures does not affect sEV yield. Size distribution was measured in NTA. Dotted lines represent standard error of the mean from 5 measurements.

Following characterization of neuron-derived EVs, we then examined whether BDNF treatment affects EV secretion. BDNF did not change the size distribution or mean particle concentration of pre-cleared supernatants as measured in NTA (**Fig. 1C**, **S1C**). Moreover, we did not observe any significant changes in the size or distribution of sEVs isolated from control (Ctrl) or BDNF-treated cortical neurons using STEM (**Fig. 1D**, **S1D-E**). Furthermore, BDNF treatment did not affect the relative abundance of TSG101 and Flotilin-2 (**Fig. 1E**, **S1F**), or the total protein and RNA concentration in cell lysates (CL) and sEVs (**Fig. 1F**, **S1G**). Overall, this data demonstrates that the majority of neuronal extracellular particles are small in size and that BDNF does not affect EV secretion.

### BDNF- but not control-stimulated sEVs increase dendrite complexity

BDNF is a well-known regulator of neuronal dendrite development, which is particularly important in the hippocampus^39–41^. To investigate whether sEVs participate in BDNF-dependent processes we first examined the effect of sEVs on dendrite complexity. Here, we incubated sEVs derived from Ctrl- or BDNF-treated cortical neurons (hereafter referred to Ctrl-EV and BDNF-EV, respectively) with hippocampal neurons at 6-7 DIV and quantified dendrite complexity three days later using Sholl analysis. Interestingly, BDNF-EVs but not Ctrl-EVs, increased dendrite complexity similarly to BDNF treatment (**Fig. 2A-C**, **S2A**). This effect was not dose-dependent, as BDNF-EVs supplied at three times the concentration had similar fold changes in complexity. Moreover, BDNF-fold changes tended to be larger and more significant in EV versus non-EV conditions, when compared to their control counterparts; Ctrl-EV or control vehicle respectively (**Fig. 2C**). To confirm that this phenotype was not mediated by residual amounts of exogenous BDNF in purified sEVs, we treated donor neurons with either recombinant BDNF, myc, or myc-BDNF and processed sEVs and CL in western blot. Neither endogenous mature BDNF, nor exogenous myc-BDNF could be detected using antibodies against BDNF or myc in sEV fractions (**Fig. S2B**). In agreement with this result, unlike BDNF treatment, incubation of BDNF-EVs with hippocampal neurons for either 20 or 60min did not lead to the phosphorylation of the BDNF receptor TrkB and did not activate downstream kinases ERK and AKT (**Fig. 2D-G**, **S2C**). Moreover, neither ERK phosphorylation nor mature BDNF expression could be detected after 3 hours of incubation with BDNF-EVs (**Fig. 2H**, **S2D-E**). Furthermore, BDNF-EVs did not induce TrkB-dependent transcriptional induction of Arc mRNA^42^ for up to 48 hours of treatment (**Fig. S2F**). Therefore, BDNF-EVs promote dendrite complexity in naïve hippocampal neurons in a mechanism that is distinct from BDNF-TrkB signaling.

**Figure 2:**
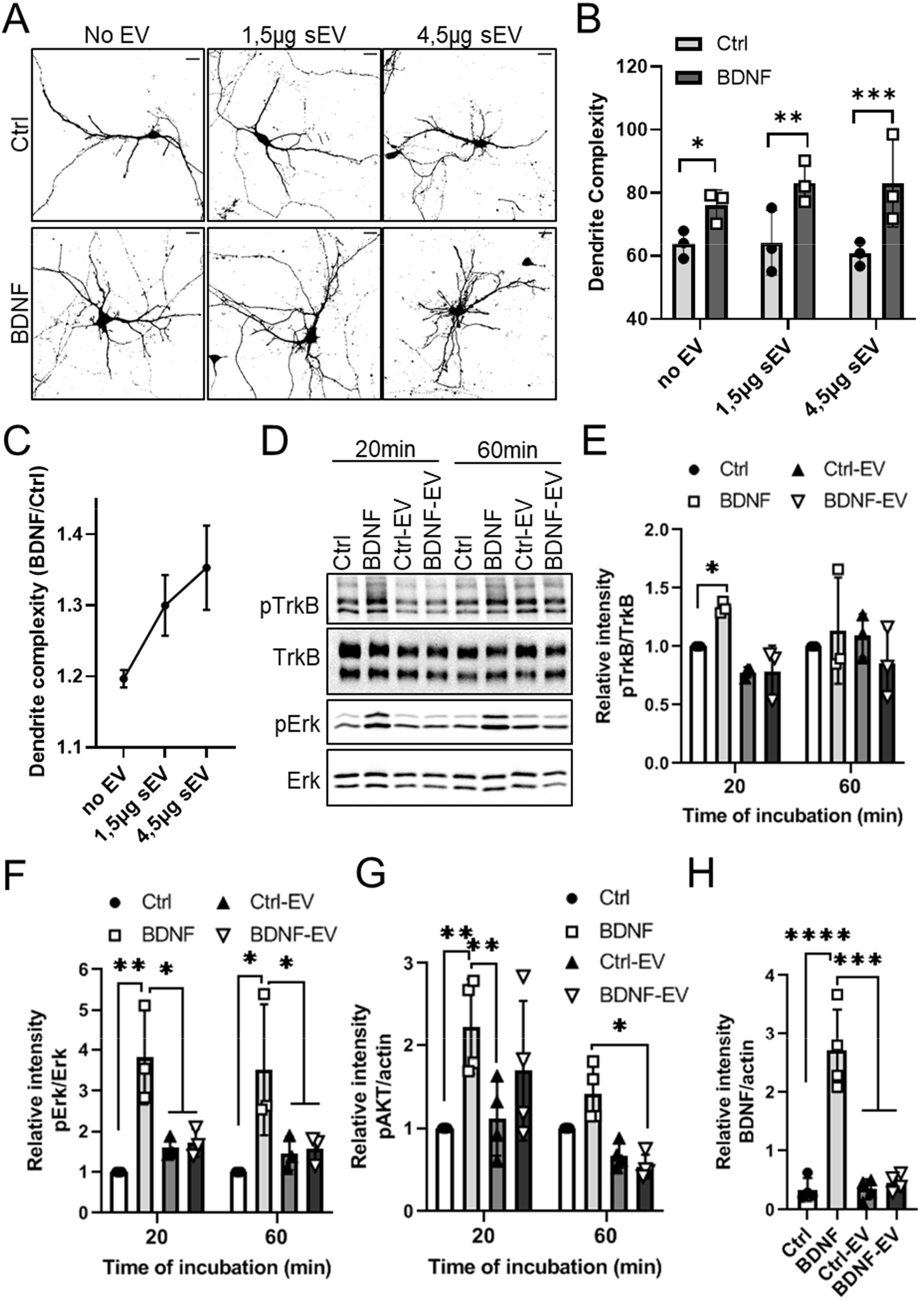
BDNF-induced sEVs increase dendrite complexity in the absence of TrkB activation and downstream signaling. **A)** BDNF-EVs increase dendrite complexity. Hippocampal neurons transfected with dsRed were treated with control vehicle (Ctrl) or BDNF (100ng/ml, 20 min) (‘No EV’), and sEVs derived from Ctrl- or BDNF-treated cortical neurons and fixed three days later. Shown are thresholded images of dsRed-expressing neurons. EVs were applied at 1,5 or 4,5 μg total protein; scale bars 20μm. **B)** Dendrite complexity of neurons treated as in **A** was calculated in Sholl analysis; n=3, 2way ANOVA, \**p*=0.04; \*\**p*=0.006; \*\*\**p*=0.002. **C)** BNDF-dependent fold changes in dendrite complexity. Dendrite complexity was normalized to respective control treatments in each independent experiment; n=4, error bars represent standard error of the mean. **D)** Western blot of BDNF-treated or EV-recipient hippocampal neurons. Recipient cells were lysed 20 or 60 min after treatment and lysates were immunoblotted with antibodies against total or phosphorylated (p) TrkB and ERK. **E)** BDNF-EVs do not stimulate TrkB phosphorylation. Relative intensities of phosphorylated or total TrkB as depicted in **A** were quantified and normalized to control treatment in each independent experiment; n=3, \**p*=0.04 in 2way ANOVA. **F)** BDNF-EVs do not stimulate ERK phosphorylation. Blots were quantified as in **B**; n=3, 2way ANOVA, \**p*<0.03, \*\**p*<0.01. **G)** BDNF-EVs do not stimulate AKT phosphorylation. Relative intensity of phosphorylated AKT (pAKT) was normalized to actin and control vehicle; n=3, 2way ANOVA, \*\**p*<0.005, \**p*=0.03. **H)** EVs do not increase mature BDNF levels. Recipient neurons were lysed after 3 hours of treatment and processed in immunoblotting. Relative intensity of mature BDNF was normalized to actin; n=3, 1way ANOVA, ***p<0.0001.

### BDNF promotes the sorting of neuronal growth-related miRNAs in sEVs

Previous publications have demonstrated the functional delivery of miRNAs between cells in several biological contexts^5,8,26,28,29^. As BDNF regulates miRNA biogenesis^33,34,38^, we were prompted to investigate whether BDNF also regulates the sorting of miRNAs into EVs. To this end, we isolated CL and corresponding sEVs from Ctrl- and BDNF-treated cortical neurons and performed next generation small RNA sequencing (NGS). Principle component analysis (PCA) revealed differential clustering of Ctrl and BDNF treated samples in both CL and sEVs (**Fig. S3A**). Interestingly however, BDNF-induced changes in individual miRNAs were not correlated between CL and sEVs (**Fig. 3A-B**). Among the most significantly changing sEV miRNAs miR-690, miR-218-5p and miR-351-5p are only regulated in sEVs, miR-132-5p is up-regulated by BDNF in both compartments and miR-129-2-3p is down-regulated in sEVs and up-regulated in CL (**Fig. 3B**, **S3B,** see also **Table S1** in *Supplementary Data*). Importantly, highly expressed sEV miRNAs, miR-132-5p, miR-218-5p, miR-690 and miR-181a-5p were largely absent from non-conditioned cell culture media (**Fig. S3D**) and with the exception of miR-690, were depleted from sEVs following inhibition of exosome secretion by GW4869 (**Fig. 3C**). Using independent qPCR experiments we further validated the BDNF-dependent up-regulation of miR-132-5p, miR-218-5p and miR-690 in sEVs, which was blocked by the specific ERK inhibitor U0126 (**Fig. 3D-F**). Importantly, neither BDNF nor U0126 treatment changed the concentration or size distribution of purified sEVs (**Fig. S3D-E**). To verify that BDNF changes sEV-associated miRNAs and not miRNAs present in the extracellular medium, we purified sEVs using size-exclusion chromatography and assessed miRNA abundance in pooled fractions enriched in either small EVs or proteins (fraction number 7-14 and 15-22 respectively). Although both fractions contained miRNAs, BDNF-induced up-regulation of miR-132-5p, miR-218-5p and miR-690 was only evident in sEV containing fractions (**Fig. 3G**). Overall, this data suggest that BDNF regulates the specific sorting of miRNAs in neuronal sEVs such as exosomes, that does not simply represent changes in intracellular miRNA abundance.

**Figure 3:**
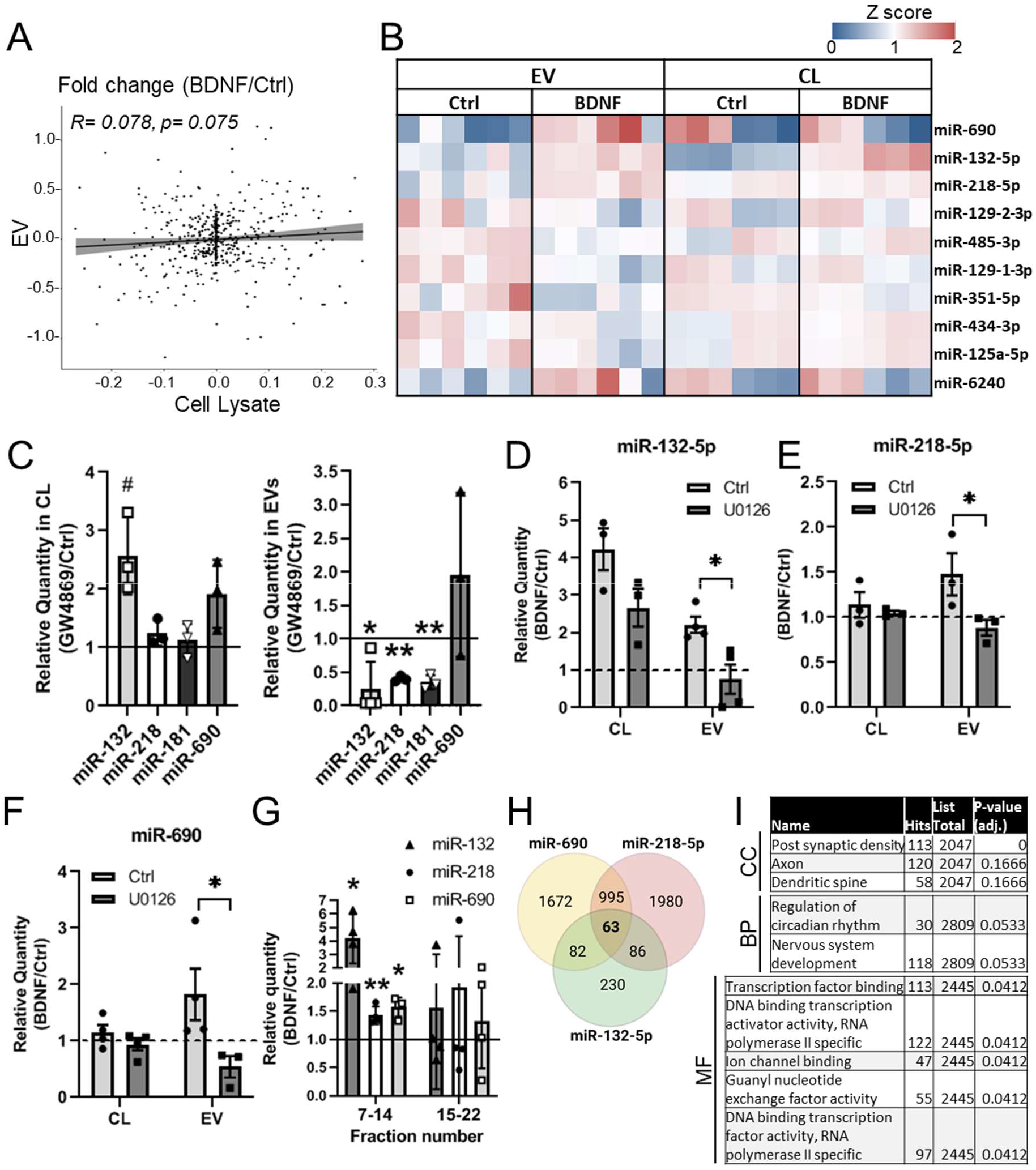
BDNF regulates the sorting of EV-miRNAs. **A)** BDNF-induced changes in miRNA abundance in EVs and cell lysates do not correlate. Depicted are Pearson’s correlation values. **B)** Heatmap depicting the levels of the top 10 regulated EV-miRNAs in decreasing order of significance and corresponding changes in cell lysates (CL); n=6. Values were normalized to the base mean for each miRNA. **C)** Inhibition of exosome secretion reduces miR-218, miR-132 and miR-181 abudnance in sEVs. Relative miRNA quantity in control or GW4869-treated cell lysates (*left*) and EVs (*right*) was quantified in rt-qPCR and normalized to RNA and control treatment; n=3-4, \**p*=0.035, \*\**p*<0.01, Student’s t-test (heteroscedastic). **D-F)** Validation of up-regulated EV-miRNAs by rt-qPCR. Treatment of EV donors with the MAPK inhibitor U0126 blocked the BDNF-induced increase in EV miRNAs. Relative values were normalized to RNA concentration and miR-181a reference miRNA; n= 3-4, \**p*<0.05, 2way ANOVA. **G)** BDNF upregulates miR-132, miR-218 and miR-690 in EV-enriched fractions isolated by size-exclusion chromatography. RNA abundance was normalized to control treatment in each independent experiment; n=4, \**p*<0.04; \*\**p*=0.006, Student’s t-test (heteroscedastic). **H)** Venn diagram depicting the number of predicted targets for up-regulated EV miRNA. Prediction was performed using miRWalk. **I)** Gene set enrichment analysis of targets common to up-regulated EV-miRNAs, using target mining (miRWalk). Shown P-values are adjusted for multiple comparisons, CC; cellular component, BP; biological pathway, MF; molecular function.

To examine the potential function of BDNF-regulated EV-miRNAs, we performed gene ontology (GO) analysis of predicted gene targets of EV-miRNAs that passed statistical significance (p<0.05) following multiple comparisons in NGS data analysis; these are miR-132-5p, miR-218-5p and miR-690 (**Table S1** in *Supplementary Data*). Target prediction revealed many overlapping targets for these miRNAs, with 63 targets common to all three miRNAs, and at least 82 targets common to any two miRNAs (**Fig. 3H**). Using target mining and gene set enrichment analysis of common targets, we observed the enrichment of genes involved in nervous system development (GO term: *Biological Pathway; BP*) and localized at synaptic compartments (GO term: *Cellular Component; CC*) (**Fig. 3I**). Moreover, we observed significant enrichment of genes implicated in transcriptional regulation of gene expression (GO term: *Molecular Function; MF*) (**Fig. 3I**). Therefore, the combined activity of these miRNAs in recipient neurons may potentially have a drastic effect in neuronal physiology that could underlie the observed growth-promoting phenotype of BDNF-EVs.

### EVs and EV-associated miRNAs are taken up by neurons

We next assessed whether EVs are taken up by neurons using the membrane binding dye lipilight (previously known as membright^43^) to fluorescently label sEVs in confocal microscopy. This dye exhibits high signal to noise ratio due to quenching by self-aggregation, and was previously verified in small EV labeling *in vivo*^44^. Indeed, our control experiments using non-conditioned cell culture media (‘No cell control’) showed negligible fluorescent signal in recipient neurons, when compared to lipilight-labeled sEVs (**Fig. 4A**), confirming that potential dye aggregates do not account for observed fluorescence signals in recipient neurons. Using immunocytochemistry, we observed the localization of lipilight-labelled Ctrl-EV and BDNF-EV in somatodendritic regions of MAP2-positive recipient hippocampal neurons (**Fig. S4A**). Moreover, the neuronal uptake of both Ctrl- and BDNF-EVs at neuronal somatodendritic compartments was completely blocked following application of the selective dynamin inhibitor dynasore (**Fig. 4B-C**), suggesting that the vast majority of neuronal sEVs are taken up via dynamin-dependent endocytosis. Consistently, using super-resolution confocal microscopy, we observed partial colocalization between lipilight-EVs and the late endosomal marker Lamp1, whereby lipilight fluorescence intensity was highest on the luminal side of Lamp1-enclosed vesicles at the neuronal soma (**Fig. S4B**). This is in line with previous reports suggesting that EVs may release their contents at late endosomal compartments^45,46^.

**Figure 4:**
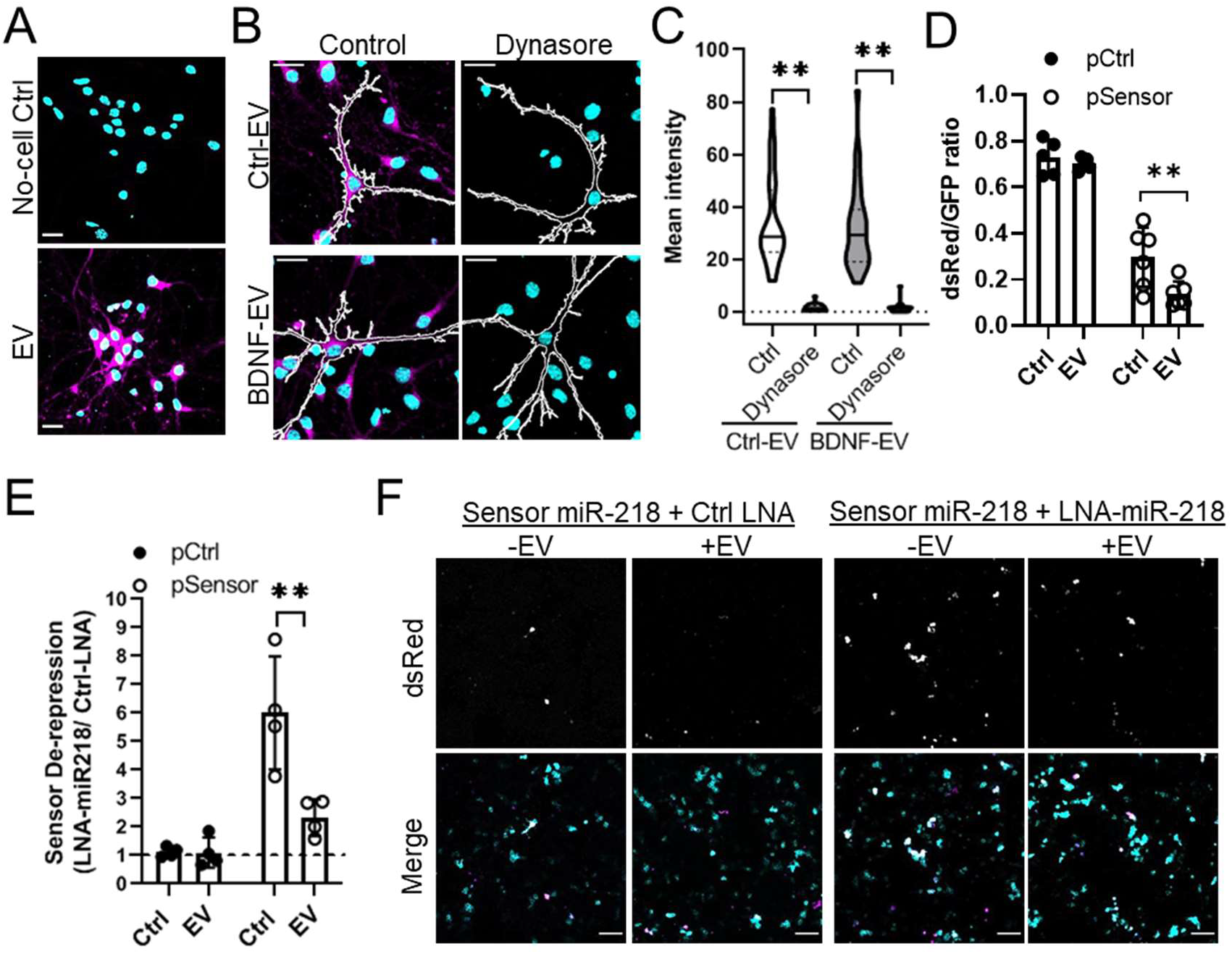
EVs and EV-miRNAs are taken up by neurons. **A)** Lipilight-560 specifically labels EVs. EVs and non-conditioned medium (No-cell Ctrl) fractions processed in parallel were labeled with lipilight-560 and added to neurons for 75 minutes. **B)** Dynasore blocks the uptake of neuronal sEVs. Hippocampal neurons pre-treated with control vehicle or Dynasore were incubated with lipilight-560-labelled sEVs for 30 minutes. White traces mark the outlines of membrane-GFP (mGFP)-expressing hippocampal neurons. Lipilight-560; magenta, DAPI; cyan, scale bars 20μm. **C)** Lipilight fluorescence intensity was quantified in the somatodendritic compartment of recipient neurons treated as in **E**. Shown is the distribution, median (black lines), upper and lower quartiles (dashed lines) from 10-15 neurons per experimental condition; n=3, 2way ANOVA, \*\**p*<0.02. **D)** N2A cell-derived sEVs increase the translational repression of a dual fluorescence reporter for miR-218 activity. Recipient N2A cells were transfected with control (pControl) and miR-218 reporter plasmid (pSensor), containing two miR-218 binding sites at the 3’UTR of dsRed and GFP as internal control. The number of dsRed-expressing and GFP-positive cells was counted approximately 20 hours following addition of EVs, n=5-6; *p=0.002, 2way ANOVA. **E)** N2A sEV supplementation reverses miR-218 sensor de-repression following inhibition of intracellular miR-218. N2A cells were co-transfected with sensor plasmids as in **D** and either Ctrl-LNA or LNA-miR-218. Sensor repression was normalized to Ctrl-LNA in each experiment; n=4, 2way ANOVA, \**p*<0.01. **F)** Representative images of dual fluorescence sensor assay as in **E**. GFP; cyan, dsRed; magenta, scale bar; 100μm.

We next examined whether our candidate miRNAs can be functional in recipient cells, using a dual fluorescence sensor plasmid for miRNA activity that consists of reporter dsRed and control GFP coding sequences downstream two separate promoters^34^. As N2a cell-derived EVs contain high levels of miR-218, we designed a sensor containing two miR-218-5p binding sites at the 3’ untranslated region (UTR) of dsRed (**Fig. S4C**). Following 20 hours of incubation, N2a cell-derived sEVs decreased the number of dsRed-expressing recipient cells transfected with the miR-218 sensor, but not control plasmid (**Fig. 4D**), suggesting that miR-218 binding is necessary for EV-mediated translational repression. Although repression of the miR-218 sensor was observed in recipient cells in the absence of sEVs, supplementation of sEVs significantly increased repression by approximately 2-fold (216.5 +/− 25.9 %). To further verify that this effect is dependent on sEV-miR-218, we transfected recipient cells with anti-sense locked nucleic acids (LNAs) to inhibit intracellular miR-218, and supplemented N2a cells with EVs 24 hours later. As expected, anti-miR-218 but not control LNAs increased the basal expression of dsRed in miR-218 sensor but not control sensor-expressing cells (**Fig. 4E**, **S4D**). This was partially rescued by sEV supplementation, which significantly decreased the number of dsRed expressing cells compared to no-EV controls (**Fig. 4E-F**). Taken together, these results demonstrate that EV-miR-218 is functional in recipient cells, and can partially rescue inhibition of intracellular miR-218 activity.

### BDNF-regulated EV miRNAs mediate dendritogenesis

To investigate whether EV miRNAs mediate the effects of BDNF-EVs on dendrite complexity, we first examined whether intracellular inhibition of BDNF-regulated EV-miRNAs, miR-218-5p, miR-132-5p and miR-690, blocks BDNF-dependent dendritogenesis. Hippocampal neurons were transfected with dsRed and either control LNA or LNA anti-sense to candidate miRNAs, after which neurons were treated with BDNF and imaged three days later (**Fig. 5A**). Dendrite complexity was assessed using Sholl analysis and fold changes in BDNF-mediated dendritogenesis were compared between each condition in neurons expressing either miR-218 or miR-132, or combinations of two or all three candidate miRNAs. We observed that inhibition of individual miRNAs did not significantly affect BDNF-induced changes in dendrite complexity, whereas simultaneous targeting of miR-132 and miR-218 completely blocked BDNF-induced dendritogenesis (**Fig. 5B**, **S5A-F**). Moreover, even though the total LNA concentration was the same for each condition (30nM), inhibition of all three miRNAs; miR-132, miR-218 and miR-690, most potently blocked BDNF-mediated dendritogenesis compared to control LNA (**Fig. 5B**, **S5A&D-F**). Furthermore, BDNF-EV supplementation of neurons expressing LNAs against all three miRNAs, blocked the LNA-induced decrease in BDNF-mediated induction of dendrite complexity (**Fig. 5C-E**). This is consistent with a role for EVs in mediating BDNF-induced dendritogenesis via the delivery of miR-132, miR-218 and miR-690.

**Figure 5:**
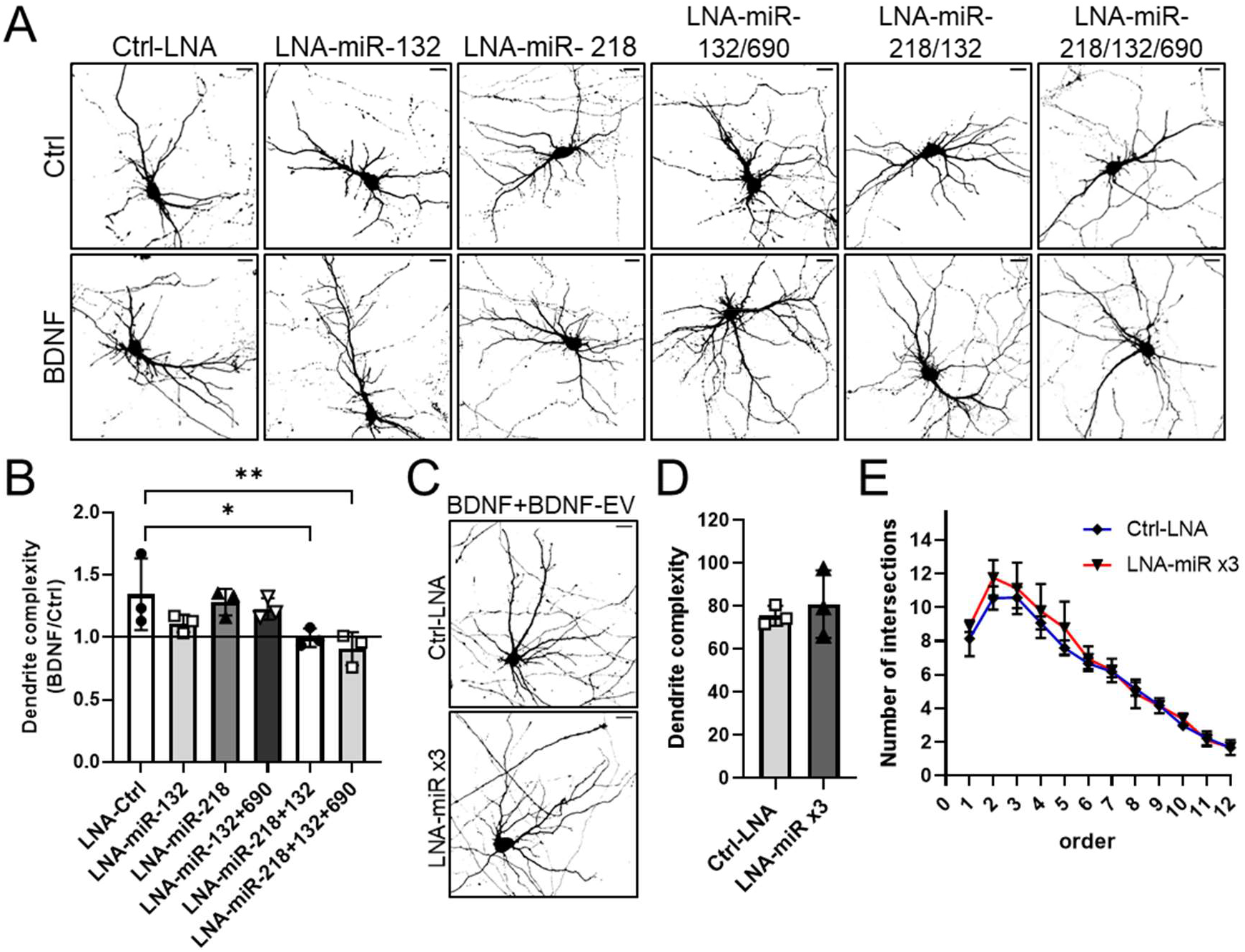
BDNF-regulated EV-miRNAs regulate dendritogenesis. **A)** Combined inhibtion of miR-128-5p, miR-132-5p and miR-690 blocks BDNF-dependent dendritogenesis. Hippocampal neurons were transfected with dsRed and control LNA (30nM), or LNAs against respective miRNAs (15nM each LNA or 10nM for co-transfection of three LNAs). Neurons were treated with BDNF (100ng/ml, 20min) and fixed three days later. Shown are thresholded images based on dsRed fluorescence, scale bars; 20μm. **B)** BDNF-fold changes in dendrite complexity were compared between each condition; n=3; 1way ANOVA; \*\**p*=0.013; \**p*=0.049. **C)** BDNF-EVs block the LNA-induced decrease in BDNF-dependent dendritogenesis. Hippocampal neurons were co-transfected with LNAs against miR-128-5p, miR-132-5p and miR-690 (‘LNA-miR x3’) and treated with BDNF as in **A**. BDNF-EVs were incubated following BDNF treatment for three days, scale bars; 20μm **D)** Dedrite complexity of neurons treated as in **C** was quantified in Sholl analysis, n=3. **E)** Sholl profiles of hippocampal neurons treated as in **C**. Error bars represent standard error, n=3.

### BDNF-EVs up-regulate synaptophysin clustering at dendrites

Our experiments suggest that BDNF-EVs promote dendrite complexity at a developmental time point corresponding to dendritic spine and synapse formation. We therefore examined whether BDNF-EVs may also influence synapse maturation. First, hippocampal neurons were treated with Ctrl- or BDNF-EVs and immunostained at 9-10DIV with antibodies against the excitatory post-synaptic marker PSD95 and the pre-synaptic marker synaptophysin (SYP) (**Fig. 6A**). Co-localization analysis of dendritic segments revealed a small but significant increase in the overlap between PSD95 and SYP in BDNF-EV compared to Ctrl-EV treated neurons (**Fig. 6B**, **S6A**), suggesting increased synapse formation. Further analysis indicated an increase in the relative intensity of SYP, but not PSD95 upon BDNF-EV treatment (**Fig. S6B**). Interestingly, this was not the case when EV donor neurons were treated with GW4869 (**Fig. 6C-D**), suggesting that exosomes are necessary for this phenotype. As SYP is a general pre-synaptic marker, we further examined whether BDNF-EVs may also affect the density of inhibitory synapses using antibodies against the post-synaptic marker Gephyrin and the pre-synaptic vesicular GABA transporter (vGAT). Here, we observed a small but significant decrease in the intensity of Gephyrin, but not vGAT at neuronal dendrites treated with BDNF-EVs compared to Ctrl-EVs (**Fig. 6E**, **S6C**). Overall this data suggest that BDNF-EVs may promote the maturation of excitatory synapses by inducing SYP clustering at dendrites.

**Figure 6:**
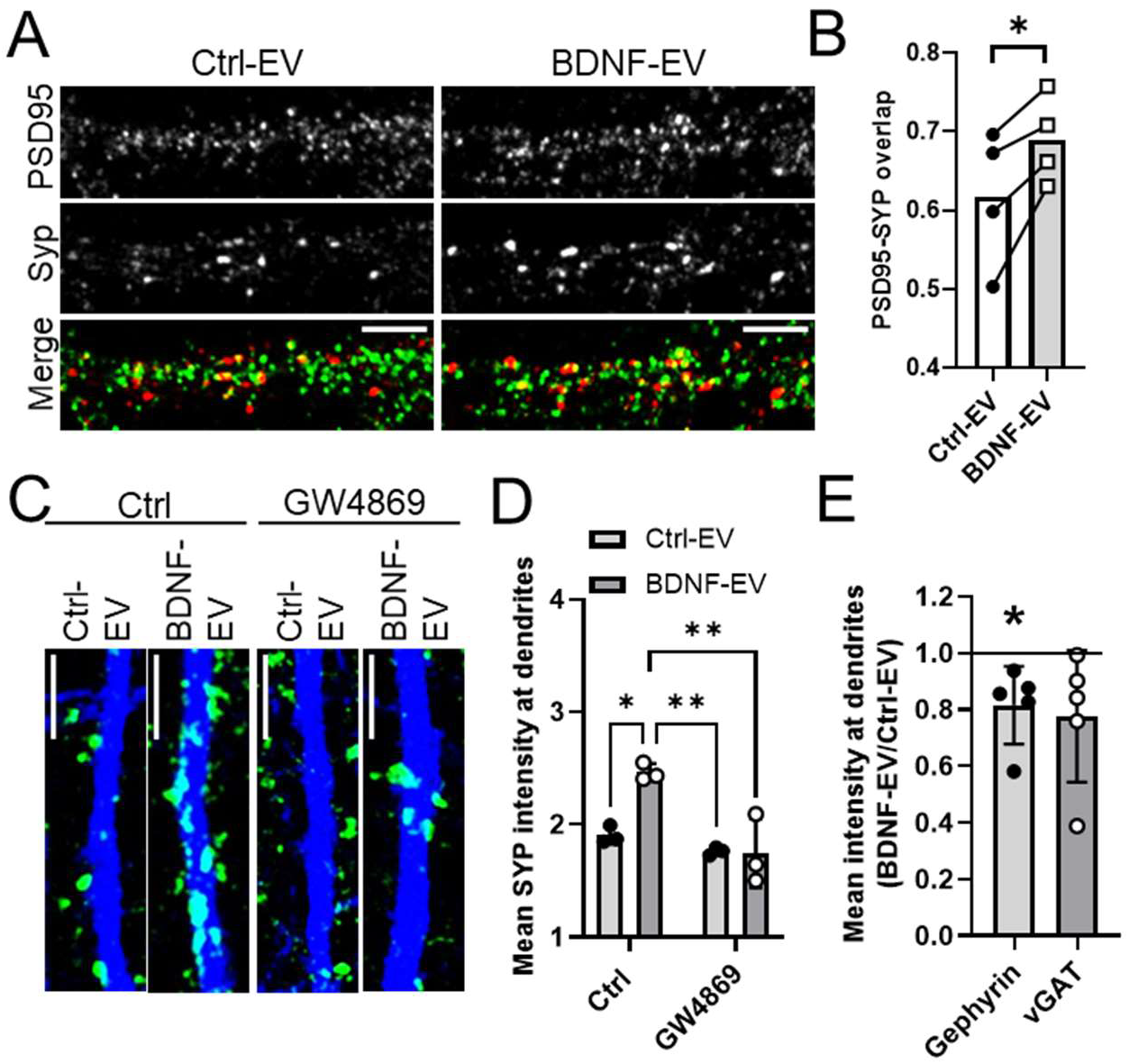
BDNF-EVs up-regulate Synaptophysin clustering at dendrites. **A)** Representative images of neuronal dendrites from EV-treated neurons. Hippocampal neurons were treated with Ctrl- or BDNF-EVs for three days and immunostained with antibodies against PSD95 (green) and synaptophysin (SYP; red). Scale bars are 5μm. **B)** BDNF-EVs increase co-localization between PSD95 and SYP. Mander’s overlap coefficients were calculated in 20μm-long dendritic segments treated as in **A**, n=4, \**p*=0.03, Student’s paired t-test. **C)** BDNF-EVs increase the levels of SYP at neuronal dendrites, which is blocked by GW4869. Ctrl- or BDNF-EVs were isolated from donor neurons treated with control vehicle or GW4869 and incubated with recipient hippocampal neurons. Recipient neurons were fixed and immunostained with anti-SYP (green) and anti-MAP2 (blue) antibodies, scale bar; 5μm. **D)** Mean fluorescence intensity of SYP was quantified in MAP2-positive neuronal dendrites as shown in **C**, n=3, 2way ANOVA, \**p*=0.01; \*\**p*<0.004. **E)** BDNF-EVs decrease gephyrin intensity in neuronal dendrites. Mean fluorescence intensity of Gephyrin and vGat was quantified in 20μm-long dendritic segments; n=5, Welch’s t-test, \**p*<0.05.

### BDNF-EVs induce synapse maturation via miRNA-218/-132/-690

We next examined whether BDNF-regulated EV-miRNA candidates are implicated in BDNF-dependent synapse maturation. Hippocampal neurons were transfected with Ctrl LNA or LNAs against the three up-regulated miRNAs; miR-218-5p, miR-132-5p and miR-690 as previously, and neurons were fixed and immunostained with PSD95 and SYP antibodies (**Fig. 7A**). Using dsRed as a morphological marker, we first examined whether dendritic spine density was affected in these conditions. Neither BDNF nor LNA-mediated inhibition of candidate miRNAs changed the number of spines along 20μm dendrites (**Fig. 7B, S6D**). We then counted the number of spines that contain PSD95 and/or are adjacent to SYP puncta. BDNF treatment alone selectively increased the number and intensity of SYP puncta opposing dendritic spines, which was not the case for PSD95 (**Fig. 7A, C**, **S6E**). Nevertheless, BDNF increased the total number of spines containing PSD95 that are adjacent to SYP suggesting that increased SYP clustering at dendrites corresponds to increased synapse formation (**Fig. 7A, D**). These effects were completely blocked by LNA-induced inhibition of BDNF-EV-regulated miRNAs (**Fig 7A, C-D**, **S6E**). Furthermore, supplementation with BDNF-EVs but not Ctrl-EVs rescued the LNA-induced decrease in SYP clustering without affecting mean PSD95 intensity (**Fig. 7E-F**) or spine density (**Fig. 6G**). Therefore, this data shows that BDNF-EVs mediate synapse maturation via BDNF-regulated miRNAs.

**Figure 7:**
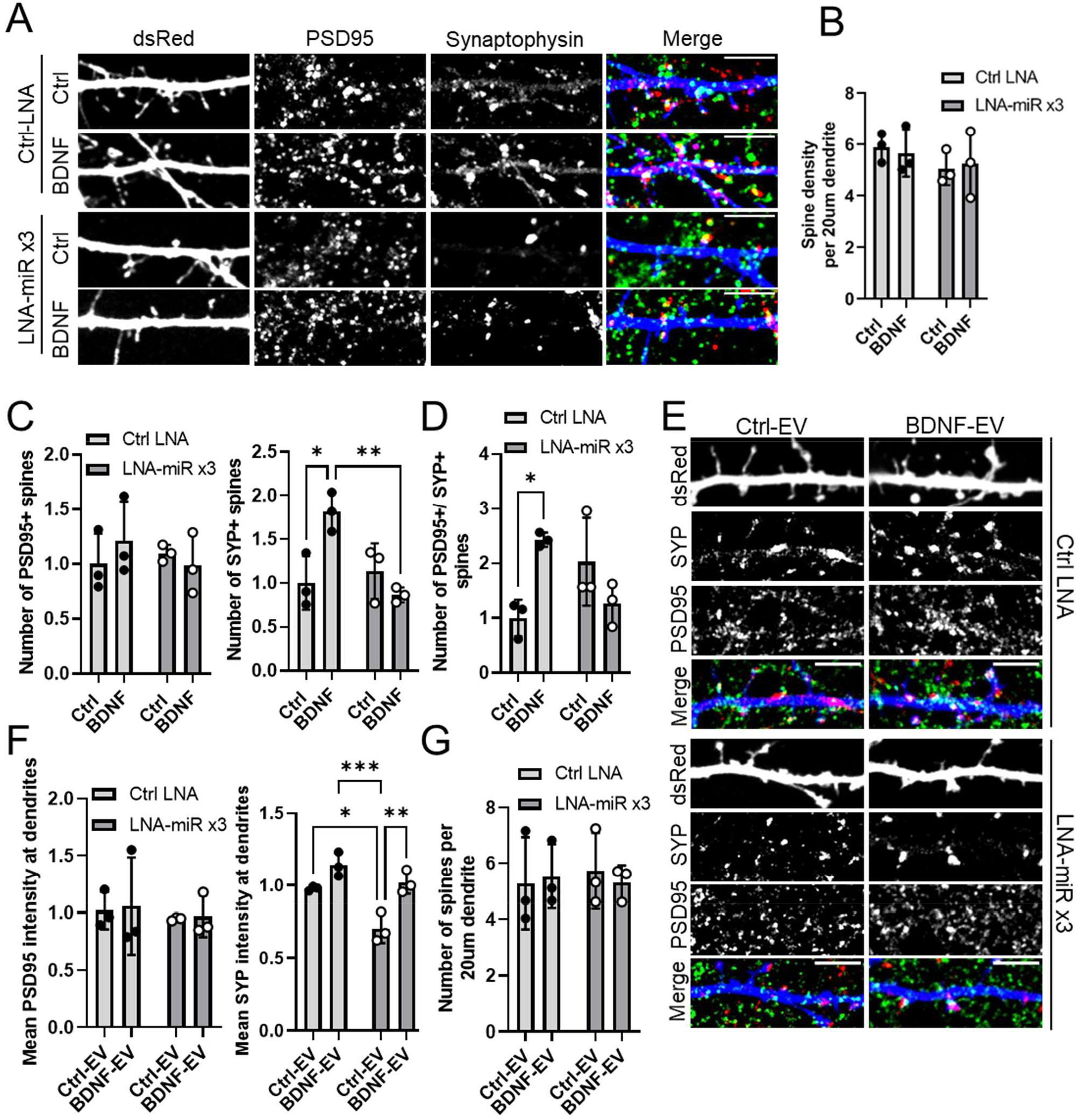
BDNF-EV up-regulation of Syp clustering is regulated via miRNAs miR-218, miR-132 and miR-690. **A)** Combined inhibition of BDNF-regulated EV-miRNAs blocks BDNF-dependent synapse maturation. Hippocampal neurons were transfected with either control LNA (30nM) or LNAs antisense to miR-218, miR-132 and miR-690 (10nM each, ‘LNA-miRx3’) and treated with BDNF (100ng/ml, 20min). Neurons were fixed three days later and immunostained with antibodies against PSD95 (green) and SYP (red); dsRed shown in blue, scale bars; 6μm. **B)** Spine density is constant across conditions. The number of dendritic spines was counted in 20μm-long, dsRed-expressing dendrites; n=3. **C)** Inhibition of BDNF-regulated EV miRNAs blocks the BDNF-induced increase in SYP-positive spines. The number of spines containing PSD95 (*left*) or SYP (*right*) was counted in 20μm-long dendrites using dsRed as a morphological marker. Values were normalized to control; n=3, 2way ANOVA, \**p*=0.02, \*\**p*=0.01. **D)** BDNF increases the number of spines containing both PSD95 and SYP, which is blocked by inhibition miR-218, miR-132 and miR-690, n=3, 2way ANOVA, \**p*=0.04. **E)** Hippocampal neurons expressing LNAs as in **A** were treated with BDNF and supplemented with Ctrl-EV or BDNF-EV. Shown are dsRed-expressing dendrites immunostained as in **A**; scale bars 5μm. **F)** BDNF-EVs rescue the LNA-induced block in BDNF-mediated SYP clustering. Mean PSD95 (*left*) and SYP (*right*) intensity was quantified in dendrites treated as in **E**, n=3, 2way ANOVA, \**p*=0.01, **p=0.004, \*\*\**p*= 0.0005. **G)** Spine density is not affected by Ctrl-EV or BDNF-EV in the presence of BDNF and LNAs, n=3.

## Discussion

We investigated the function of neuronal sEVs within the context of a well-established paradigm of neuronal morphogenesis downstream BDNF signaling. BDNF selectively regulated the sorting of growth-related miRNAs in neuronal sEVs, the majority of which did not change in cell lysates. EVs from BDNF- but not control-treated neurons induced dendrite complexity and synapse maturation in naïve hippocampal dendrites, similarly to BDNF treatment itself. Remarkably, this was not due to the activation of BDNF-TrkB signaling cascades or associated changes in transcriptional regulation. Rather, three miRNAs that were significantly up-regulated in sEV fractions, miR-132, miR-218 and miR-690, mediated the observed BDNF-EV-induced phenotypes. Therefore, our study uncovers a novel mechanism of neuronal dendrite maturation downstream BDNF and provides primary evidence of functional miRNA transfer between neurons via sEVs.

Several studies have now shown that EVs are heterogeneous, with sEV preparations likely consisting of many different EV sub-types^20,21^. As efficient separation of these sub-types is not possible so far, we refrain from further classifying sEVs in this study. Nevertheless, we observed an approximate 1.5-fold reduction of particles in sEV fractions following treatment of donor neurons with the N-SMase inhibitor GW4869, which is necessary for ESCRT-independent biogenesis of intra-luminal vesicles in MVEs destined to be secreted as exosomes^47^. Additionally, we observed the depletion of several miRNAs in sEVs derived from GW4869-treated neurons, and GW4869 treatment of EV donors abolished the BDNF-EV-dependent increase in SYP clustering. Notably, miR-690, which was regulated at vesicle fractions isolated by size-exclusion chromatography, was not depleted by GW4869, suggesting that both exosomes and non-exosomal vesicles are likely to contribute to BDNF-EV phenotypes.

Despite several examples of functional miRNA transfer between cells^5,8,9,28,29^, there is still some controversy regarding the presence of EV miRNAs at sufficient concentrations^48^, and the efficiency of EV uptake by the recipient cells^46^. Admittedly however, these studies do not exclude the possibility that these processes may become more favorable under specific conditions, for instance based on the type and state of donor and recipient cells. Our results using the single cell reporter for miRNA activity indeed demonstrate that EV miR-218 is functional and administration of EVs largely rescues decreased reporter repression following inhibition of intracellular miR-218 in recipient N2A cells. Moreover, we show that the co-targeting of at least two EV-miRNAs is necessary for a complete block in BDNF-EV phenotypes. As these miRNA candidates share many common targets, which are predicted to be involved in neurodevelopmental processes, a co-targeting mechanism may provide a competitive advantage for EV-miRNA-mediated translational repression. In this scenario, binding of multiple miRNA-induced silencing complexes (miRISC) to a 3’UTR would have a stronger effect on translational repression, presumably via more efficient recruitment of associated factors involved in the silencing or degradation of the target mRNA. This effect was recently demonstrated experimentally for co-targeting of several neuronal transcripts by two or more miRNAs^49^, and is also supported by studies showing that specific groups of miRNAs are necessary for a biological response such as neuronal differentiation^50,51^.

Notably, miRNAs that were up-regulated in BDNF-EVs were previously implicated in BDNF-related neuronal processes. For instance, miR-132-5p is part of the miR-132/-212 gene cluster that is transcriptionally induced by BDNF^35^. Deletion or overexpression of these miRNAs led to defects in memory and hippocampal plasticity. Although most publications define a function for miR-132-3p, which has a different set of targets than miR-132-5p, so far the relative contribution of each mature miRNA from this cluster is unclear^52^. Interestingly, we could only validate the regulation of miR-132-5p in BDNF-EVs, whereas other members of the cluster were not affected. It is therefore intriguing to speculate that miR-132-5p is selectively sorted in EVs via an undefined mechanism, which could explain the reported low amounts in cells. The two other miRNAs miR-218-5p and miR-690, were selectively up-regulated in sEVs and not CL, and were the only arms that were detectable by NGS in either compartment. In line with a role in BDNF-dependent processes suggested by our results, miR-218 is enriched in neurites and was previously shown to promote increased synaptic strength ^53^, as well as resistance to stress-induced depressive behaviors^54^. The latter was positively correlated to peripheral miR-218 levels. Although a potential role for EVs was not addressed in this study, EVs from the brain are found in the blood and cerebrospinal fluid, and EV-miRNAs are a promising source of biomarkers of cognitive decline and early prognosis of diseases like Alzheimer’s disease^55,56^.

Interestingly, whereas BDNF signaling itself is thought to be highly localized to sites of BDNF secretion, reportedly only up to 4-5 microns^61^, several examples have demonstrated that EVs are capable of traveling long distances, for instance crossing tissue barriers such as the placenta^57^ and the blood brain barrier^58^, and across synapses^10,59,60^. Moreover, we previously suggested the localized regulation of miRNA production by BDNF via the dynamic anchoring of the miRNA production machinery at membranes of dendritic organelles^34^. Although further work is needed to elucidate the precise mechanisms of EV-miRNA sorting, secretion and potential spreading, localized BDNF-EV secretion may occur within the vicinity of TrkB activation sites. This may be important for regulating the morphology of neighboring or inter-connected neurons and potentially prime synapses for subsequent responses to increased neuronal activity. Given the wide-spread role of BDNF signaling in hippocampal plasticity, EVs may thus contribute to neurobiological processes underlying learning and memory.

## Supplementary Files

Supplementary Methods

- List of Materials Tables S1&S2
- Supplementary Methods

Supplementary Data

- Supplementary Figures S1-S6
- Supplementary Data Table S1

## Acknowledgements

We thank Ms Julia Lindlar for technical help with EV isolation. We are grateful to Dr. Melania Capasso and Prof. Donato Di Monte for kindly sharing antibodies. Prof. Susanne Schoch and Prof. Gerhard Schratt generously provided DNA plasmids. We thank the DZNE Light Microscope core facility (LMF) for providing support and instrumentation for light microscopy experiments and the Microscopy Core Facility of the Medical Faculty at the University of Bonn for their support in scanning transmission electron microscopy, funded by the German Research Foundation (Deutsche Forschungsgemeinschaft, DFG – Projectnumber 388171357). AA was funded via the BONFOR research foundation of the Bonn Medical Faculty for implementation of this project.

## Author declaration

All authors have seen and approved this manuscript and confirm that it has not been accepted for publication elsewhere.

## References

1. van Niel, G., D’Angelo, G. & Raposo, G. Shedding light on the cell biology of extracellular vesicles. Nat. Rev. Mol. Cell Biol. 19, 213–228 (2018).

2. Frühbeis, C. et al. Neurotransmitter-triggered transfer of exosomes mediates oligodendrocyte-neuron communication. PLoS Biol. 11, e1001604 (2013).

3. Mukherjee, C. et al. Oligodendrocytes provide antioxidant defense function for neurons by secreting ferritin heavy chain. Cell Metab. 32, 259–272.e10 (2020).

4. Chaudhuri, A. D. et al. TNFα and IL-1β modify the miRNA cargo of astrocyte shed extracellular vesicles to regulate neurotrophic signaling in neurons. Cell Death Dis. 9, 363 (2018).

5. Men, Y. et al. Exosome reporter mice reveal the involvement of exosomes in mediating neuron to astroglia communication in the CNS. Nat. Commun. 10, 4136 (2019).

6. Bahrini, I., Song, J., Diez, D. & Hanayama, R. Neuronal exosomes facilitate synaptic pruning by up-regulating complement factors in microglia. Sci. Rep. 5, 7989 (2015).

7. Glebov, K. et al. Serotonin stimulates secretion of exosomes from microglia cells. Glia 63, 626–634 (2015).

8. Prada, I. et al. Glia-to-neuron transfer of miRNAs via extracellular vesicles: a new mechanism underlying inflammation-induced synaptic alterations. Acta Neuropathol. 135, 529–550 (2018).

9. Xu, B. et al. Neurons secrete miR-132-containing exosomes to regulate brain vascular integrity. Cell Res. 27, 882–897 (2017).

10. Wang, Y. et al. The release and trans-synaptic transmission of Tau via exosomes. Mol. Neurodegener. 12, 5 (2017).

11. Asai, H. et al. Depletion of microglia and inhibition of exosome synthesis halt tau propagation. Nat. Neurosci. 18, 1584–1593 (2015).

12. Wang, B. & Han, S. Exosome-associated tau exacerbates brain functional impairments induced by traumatic brain injury in mice. Mol. Cell. Neurosci. 88, 158–166 (2018).

13. Sardar Sinha, M. et al. Alzheimer’s disease pathology propagation by exosomes containing toxic amyloid-beta oligomers. Acta Neuropathol. 136, 41–56 (2018).

14. Lim, C. Z. J. et al. Subtyping of circulating exosome-bound amyloid β reflects brain plaque deposition. Nat. Commun. 10, 1144 (2019).

15. Stuendl, A. et al. Induction of α-synuclein aggregate formation by CSF exosomes from patients with Parkinson’s disease and dementia with Lewy bodies. Brain 139, 481–494 (2016).

16. Xia, Y. et al. Microglia as modulators of exosomal alpha-synuclein transmission. Cell Death Dis. 10, 174 (2019).

17. Sharma, P. et al. Exosomes regulate neurogenesis and circuit assembly. Proc. Natl. Acad. Sci. USA 116, 16086–16094 (2019).

18. Lee, S. H. et al. Reciprocal control of excitatory synapse numbers by Wnt and Wnt inhibitor PRR7 secreted on exosomes. Nat. Commun. 9, 3434 (2018).

19. Vilcaes, A. A., Chanaday, N. L. & Kavalali, E. T. Interneuronal exchange and functional integration of synaptobrevin via extracellular vesicles. Neuron 109, 971–983.e5 (2021).

20. Jeppesen, D. K. et al. Reassessment of exosome composition. Cell 177, 428–445.e18 (2019).

21. Mathieu, M., Martin-Jaular, L., Lavieu, G. & Théry, C. Specificities of secretion and uptake of exosomes and other extracellular vesicles for cell-to-cell communication. Nat. Cell Biol. 21, 9–17 (2019).

22. Valadi, H. et al. Exosome-mediated transfer of mRNAs and microRNAs is a novel mechanism of genetic exchange between cells. Nat. Cell Biol. 9, 654–659 (2007).

23. Weiss, K., Antoniou, A. & Schratt, G. Non-coding mechanisms of local mRNA translation in neuronal dendrites. Eur. J. Cell Biol. 94, 363–367 (2015).

24. Rajman, M. & Schratt, G. MicroRNAs in neural development: from master regulators to fine-tuners. Development 144, 2310–2322 (2017).

25. Olde Loohuis, N. F. M. et al. MicroRNA networks direct neuronal development and plasticity. Cell Mol. Life Sci. 69, 89–102 (2012).

26. Goldie, B. J. et al. Activity-associated miRNA are packaged in Map1b-enriched exosomes released from depolarized neurons. Nucleic Acids Res. 42, 9195–9208 (2014).

27. Li, H., Wu, C., Aramayo, R., Sachs, M. S. & Harlow, M. L. Synaptic vesicles contain small ribonucleic acids (sRNAs) including transfer RNA fragments (trfRNA) and microRNAs (miRNA). Sci. Rep. 5, 14918 (2015).

28. Das, S. & Halushka, M. K. Extracellular vesicle microRNA transfer in cardiovascular disease. Cardiovasc Pathol 24, 199–206 (2015).

29. Chen, J., Hu, C. & Pan, P. Extracellular vesicle microrna transfer in lung diseases. Front. Physiol. 8, 1028 (2017).

30. Balusu, S. et al. Identification of a novel mechanism of blood-brain communication during peripheral inflammation via choroid plexus-derived extracellular vesicles. EMBO Mol. Med. 8, 1162–1183 (2016).

31. Leal, G., Afonso, P. M., Salazar, I. L. & Duarte, C. B. Regulation of hippocampal synaptic plasticity by BDNF. Brain Res. 1621, 82–101 (2015).

32. Aakalu, G., Smith, W. B., Nguyen, N., Jiang, C. & Schuman, E. M. Dynamic visualization of local protein synthesis in hippocampal neurons. Neuron 30, 489–502 (2001).

33. Huang, Y.-W. A., Ruiz, C. R., Eyler, E. C. H., Lin, K. & Meffert, M. K. Dual regulation of miRNA biogenesis generates target specificity in neurotrophin-induced protein synthesis. Cell 148, 933–946 (2012).

34. Antoniou, A. et al. The dynamic recruitment of TRBP to neuronal membranes mediates dendritogenesis during development. EMBO Rep. 19, (2018).

35. Vo, N. et al. A cAMP-response element binding protein-induced microRNA regulates neuronal morphogenesis. Proc. Natl. Acad. Sci. USA 102, 16426–16431 (2005).

36. Schratt, G. M. et al. A brain-specific microRNA regulates dendritic spine development. Nature 439, 283–289 (2006).

37. Gao, J. et al. A novel pathway regulates memory and plasticity via SIRT1 and miR-134. Nature 466, 1105–1109 (2010).

38. Fiore, R. et al. Mef2-mediated transcription of the miR379-410 cluster regulates activity-dependent dendritogenesis by fine-tuning Pumilio2 protein levels. EMBO J. 28, 697–710 (2009).

39. Novkovic, T., Mittmann, T. & Manahan-Vaughan, D. BDNF contributes to the facilitation of hippocampal synaptic plasticity and learning enabled by environmental enrichment. Hippocampus 25, 1–15 (2015).

40. Leal, G., Bramham, C. R. & Duarte, C. B. BDNF and hippocampal synaptic plasticity. Vitam Horm 104, 153–195 (2017).

41. Heldt, S. A., Stanek, L., Chhatwal, J. P. & Ressler, K. J. Hippocampus-specific deletion of BDNF in adult mice impairs spatial memory and extinction of aversive memories. Mol. Psychiatry 12, 656–670 (2007).

42. Ying, S.-W. et al. Brain-derived neurotrophic factor induces long-term potentiation in intact adult hippocampus: requirement for ERK activation coupled to CREB and upregulation of Arc synthesis. J. Neurosci. 22, 1532–1540 (2002).

43. Collot, M. et al. Membright: A family of fluorescent membrane probes for advanced cellular imaging and neuroscience. Cell Chem. Biol. 26, 600–614.e7 (2019).

44. Hyenne, V., Lefebvre, O. & Goetz, J. G. Going live with tumor exosomes and microvesicles. Cell Adh Migr 11, 173–186 (2017).

45. Joshi, B. S., de Beer, M. A., Giepmans, B. N. G. & Zuhorn, I. S. Endocytosis of Extracellular Vesicles and Release of Their Cargo from Endosomes. ACS Nano 14, 4444–4455 (2020).

46. Bonsergent, E. et al. Quantitative characterization of extracellular vesicle uptake and content delivery within mammalian cells. Nat. Commun. 12, 1864 (2021).

47. Catalano, M. & O’Driscoll, L. Inhibiting extracellular vesicles formation and release: a review of EV inhibitors. J Extracell Vesicles 9, 1703244 (2020).

48. Chevillet, J. R. et al. Quantitative and stoichiometric analysis of the microRNA content of exosomes. Proc. Natl. Acad. Sci. USA 111, 14888–14893 (2014).

49. Cherone, J. M., Jorgji, V. & Burge, C. B. Cotargeting among microRNAs in the brain. Genome Res. 29, 1791–1804 (2019).

50. Santos, M. C. T. et al. miR-124, −128, and −137 Orchestrate Neural Differentiation by Acting on Overlapping Gene Sets Containing a Highly Connected Transcription Factor Network. Stem Cells 34, 220–232 (2016).

51. Pons-Espinal, M. et al. Synergic Functions of miRNAs Determine Neuronal Fate of Adult Neural Stem Cells. Stem Cell Rep. 8, 1046–1061 (2017).

52. Aten, S., Hansen, K. F., Hoyt, K. R. & Obrietan, K. The miR-132/212 locus: a complex regulator of neuronal plasticity, gene expression and cognition. RNA Dis. 3, (2016).

53. Rocchi, A. et al. Neurite-Enriched MicroRNA-218 Stimulates Translation of the GluA2 Subunit and Increases Excitatory Synaptic Strength. Mol. Neurobiol. 56, 5701–5714 (2019).

54. Torres-Berrío, A. et al. MiR-218: a molecular switch and potential biomarker of susceptibility to stress. Mol. Psychiatry 25, 951–964 (2020).

55. Cheng, L. et al. Prognostic serum miRNA biomarkers associated with Alzheimer’s disease shows concordance with neuropsychological and neuroimaging assessment. Mol. Psychiatry 20, 1188–1196 (2015).

56. Jain, G. et al. A combined miRNA-piRNA signature to detect Alzheimer’s disease. Transl. Psychiatry 9, 250 (2019).

57. Tannetta, D., Dragovic, R., Alyahyaei, Z. & Southcombe, J. Extracellular vesicles and reproduction-promotion of successful pregnancy. Cell Mol Immunol 11, 548–563 (2014).

58. Morad, G. et al. Tumor-Derived Extracellular Vesicles Breach the Intact Blood-Brain Barrier via Transcytosis. ACS Nano 13, 13853–13865 (2019).

59. Korkut, C. et al. Regulation of postsynaptic retrograde signaling by presynaptic exosome release. Neuron 77, 1039–1046 (2013).

60. Pastuzyn, E. D. et al. The Neuronal Gene Arc Encodes a Repurposed Retrotransposon Gag Protein that Mediates Intercellular RNA Transfer. Cell 172, 275–288.e18 (2018).

61. Horch, H. W. & Katz, L. C. BDNF release from single cells elicits local dendritic growth in nearby neurons. Nat. Neurosci. 5, 1177–1184 (2002).

