## Supplementary Data for "Neuronal extracellular vesicles mediate BDNF-dependent dendritogenesis and synapse maturation via microRNAs"

### Supplementary Figures

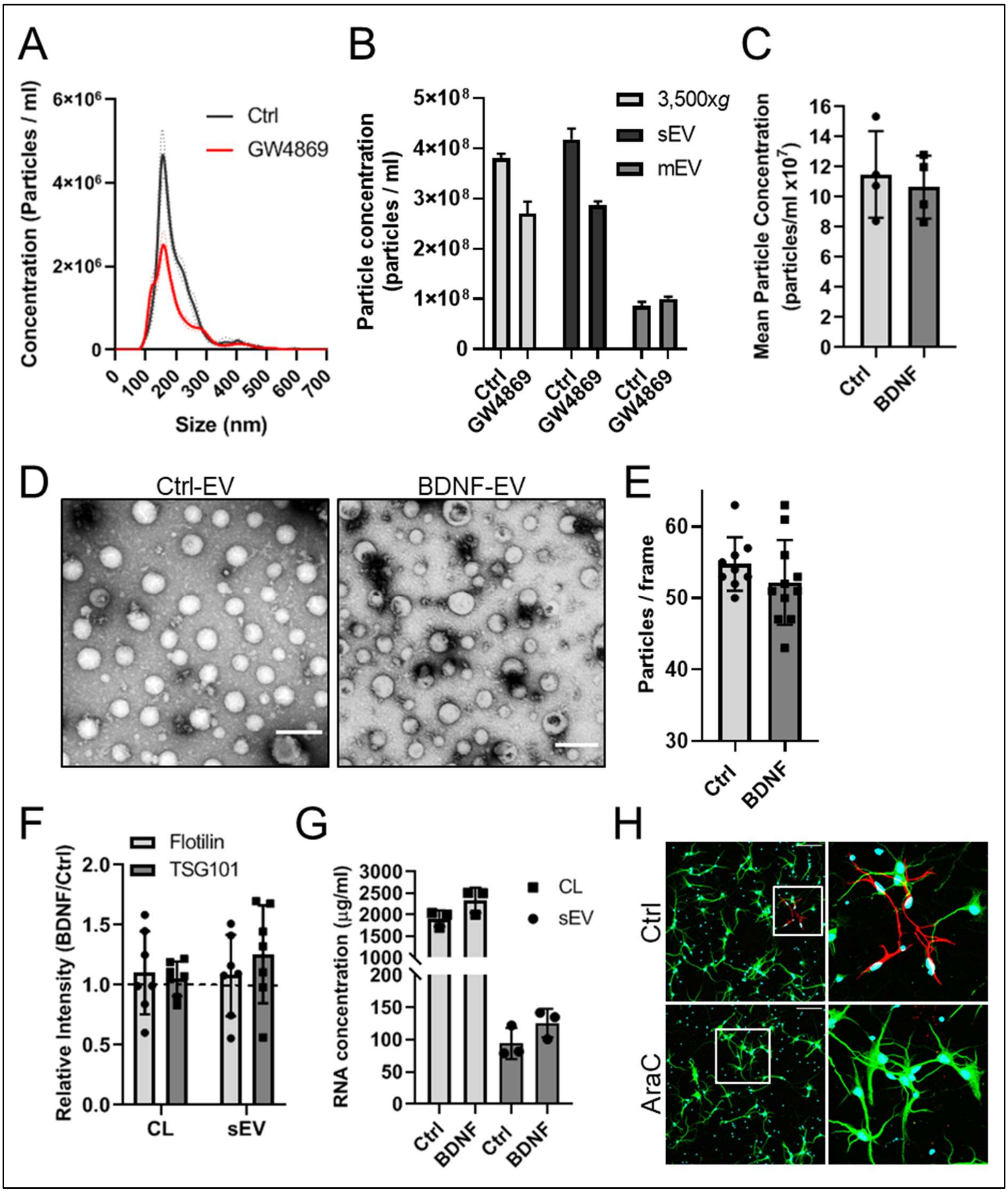


**Figure S1: BDNF does not affect EV secretion**

**A)** GW4869 reduces sEV secretion. Particle size distribution of sEVs was measured using NTA in neurons treated with control DMSO or GW4869.

**B)** Particle concentration in neuronal culture supernatants (3,500xg), sEVs and mEVs was quantified in NTA following treatment with control DMSO or GW4869. Values were normalized to protein concentration in corresponding cell lysates and dilution factor. Shown is the average and standard error of five consecutive measurements.

**C)** BDNF does not change the concentration of extracellular particles. Pre-cleared supernatants were processed in NTA in four independent experiments; n=4 (related to **Fig. 1C**).

**D)** Representative STEM images of Ctrl and BDNF-induced sEVs; scale bar is 200nm (related to **Fig. 1D**).

**E)** Particle number of sEVs imaged using STEM as shown in **B** was calculated in approximately 10 STEM micrographs for each condition (dimensions 1,14 $\mu m^2$).

**F)** Relative intensities of TSG101 and Flotilin-2 from neuronal cell lysates and corresponding sEVs treated with BDNF were normalized to control treatment in each independent experiment; n=7 (related to **Fig. 1E).**

**G)** BDNF does not change total RNA concentration in cortical neuron lysates (CL) or corresponding sEVs isolated by UC; n=3.

**H)** Representative images of control (Ctrl) or AraC-treated donor neurons immunostained with MAP2 (green) and GFAP (red); DAPI stained nucei shown in cyan, scale bar; 100µm (related to **Fig. 1G**).

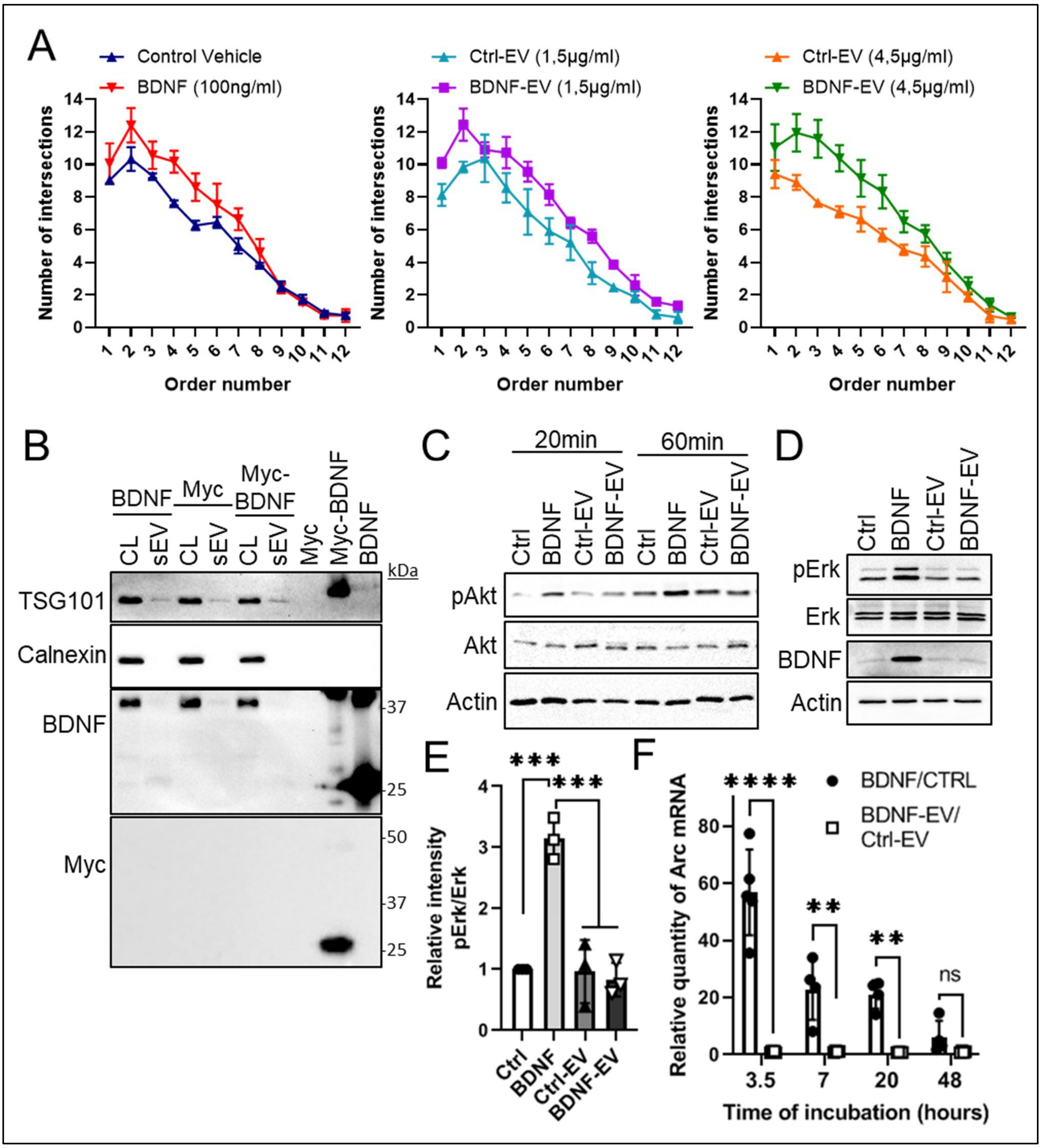


**Figure S2: BDNF-induced sEVs increase dendrite complexity in the absence of TrkB activation and downstream signaling**

**A)** Sholl profiles of hippocampal neurons treated with control, BDNF, Ctrl-EV and BDNF-EVs, as indicated. Shown are the number of intersections at each concentric circle at increasing distance from the soma. Error bars represent standard error of the mean, n=3 (related to **Fig. 2A-B**).

**B)** BDNF is not present in sEV preparations. Neurons were treated with either BDNF, myc or myc-BDNF (50ng/ml, 30min) and cell lysates (CL) and purified sEVs were collected as previously.

CL, sEVs and 50ng purified proteins; myc, myc-BDNF and BDNF, were processed in western blotting.

**C)** BDNF-EVs do not induce Akt phosphorylation (related to **Fig. 2G**)

**D)** Immunoblot of recipient neuron lysates following 3 hours of treatment. Western blot bands correspond to phosphorylated (p) and total ERK, mature BDNF and actin (related to **Fig. 2H**).

**E)** EVs do not increase ERK phosphorylation after 3 hours of incubation. Relative intensity of pErk and Erk immunoblots as shown in **B** was quantified and normalized to control treatment; n=3, 1way ANOVA, ***$p$<0.001.

**F)** BDNF-EVs do not induce mature Arc mRNA expression. Hippocampal neurons were treated with Ctrl, BDNF, Ctrl-EV or BDNF-EV and fold changes in Arc mRNA quantity were quantified by qPCR at indicated time points. Values were normalized to RNA concentration and GAPDH reference gene expression in each experiment; n=5, 2way ANOVA, **$p$<0.01, ***$p$<0.0001.

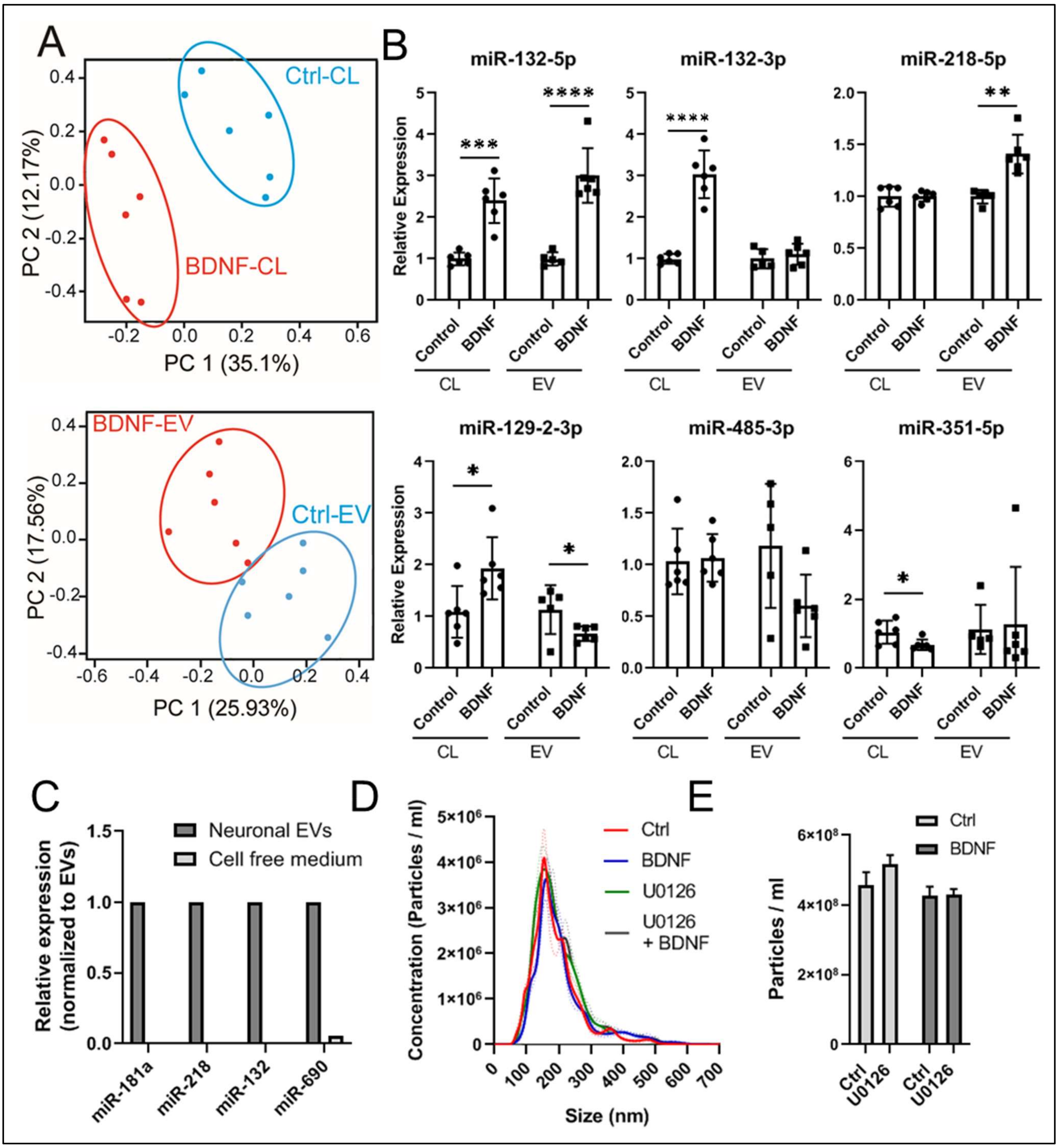


**Figure S3: BDNF regulates the sorting of EV-miRNAs**

**A)** Principle component (PC) analysis of NGS data from Ctrl or BDNF-treated cell lysates (*top*) and EVs (*bottom*).

**B)** Validation of small RNA sequencing data. MiRNA abundance was normalized to the small nuclear RNA (snRNA) U6, n=5-6, Student's t-test, ****p<0.0001; ***p=0.0001; **p=0.001; *p<0.05.

**C)** Cell-free neuronal culture media has very little or no miRNA candidates. Conditioned and non-conditioned media were processed in parallel and miRNA abundance was normalized to neuronal EV abundance for each miRNA.

**D)** BDNF and/or U0126 treatment does not change the distribution of sEVs measured by NTA. Traces and dotted lines represent the mean and standard error from five measurements (related to **Fig. 3D-F**).

**E)** Particle concentration of sEVs treated as in **D** was quantified in NTA. Shown is the average and standard error of five consecutive measurements.

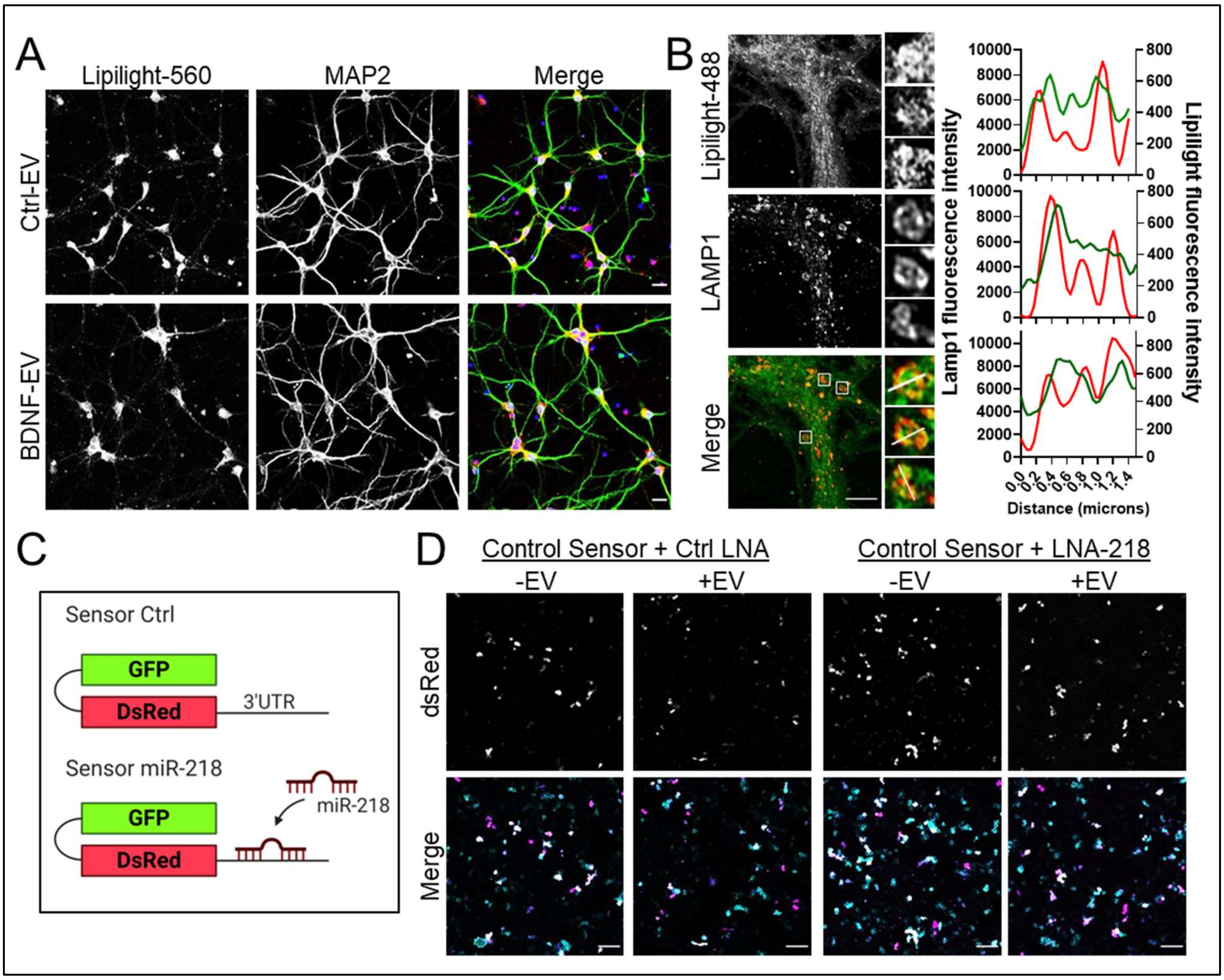


**Figure S4: EVs and EV-miRNAs are internalized by neurons**

**A)** Neurons internalize control and BDNF-stimulated sEVs at the somatodendritic compartment. Lipilight-560-labeled sEVs (red) were incubated with 7DIV hippocampal neurons for 75 minutes prior to washing and immunostaining with the neuronal marker MAP2 (green). Scale bars; 20µm, DAPI-stained nuclei shown in blue.

**B)** Internalized EVs co-localize with the late endosomal marker Lamp1. Hippocampal neurons were incubated with neuronal EVs for 75 min and immunostained for Lamp1. Image stacks were acquired using the Airy scan module. (*Right*) Line fluorescence intensity profiles are shown for each region of interest (ROI; 1.8x1.8µm) as depicted by magnified insets and white lines in merged images, EV; green, Lamp1; red, scale bar; 5µm.

**C)** Diagram depicting dual-fluorescence miRNA sensor plasmids. The miR-218 sensor contains two binding sites for miR-218-5p at the 3'untranslated region (3'UTR) of dsRed.

**D)** Representative images of dual-fluorescence sensor assay control plasmid (related to **Fig. 4E**).

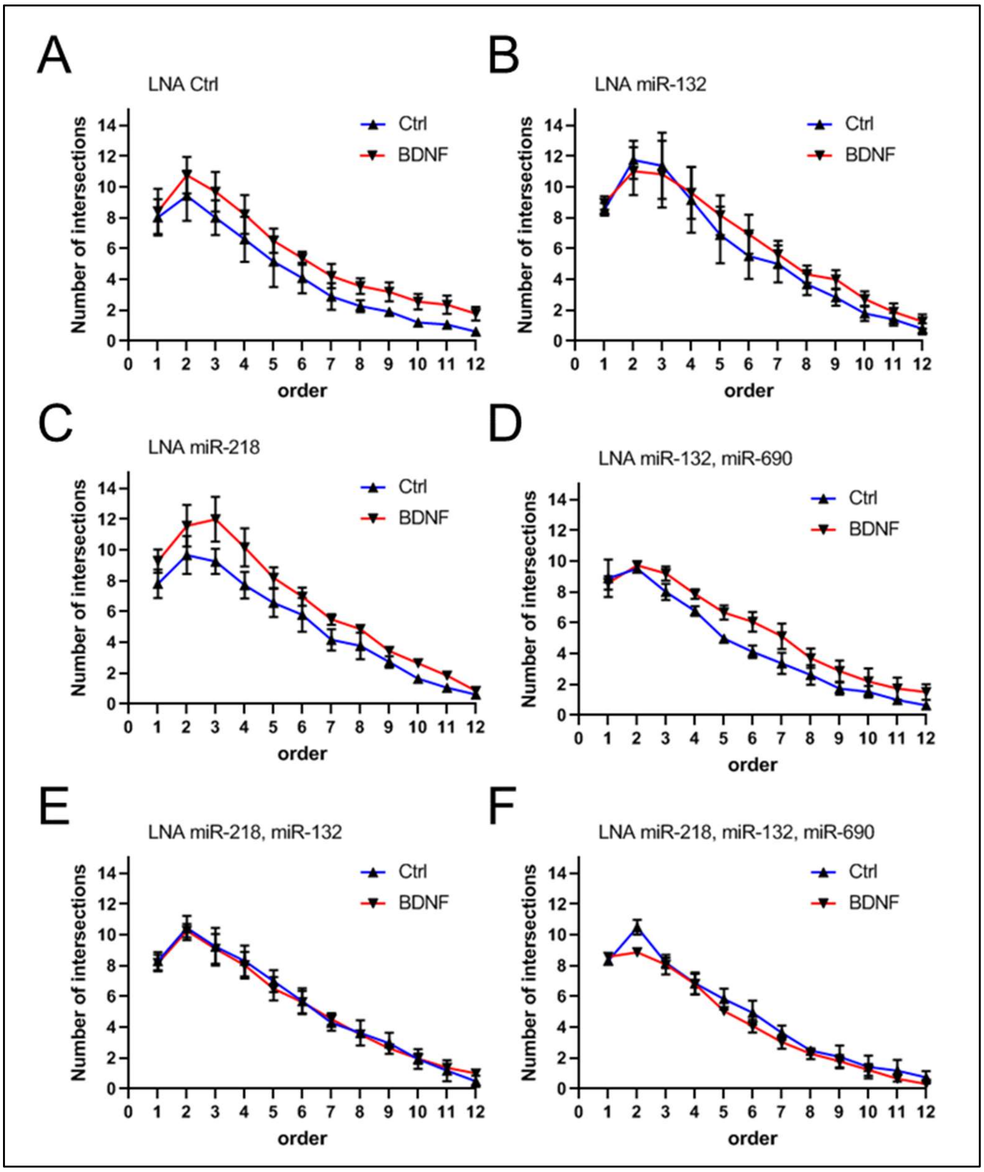


<u>**Figure S5:**</u> **BDNF-regulated EV-miRNAs regulate dendritogenesis**. **A-F)** Sholl analysis profiles are shown for each condition as depicted; n=3-4, error barrs represent standard deviation (related to **Fig. 5A-B**).

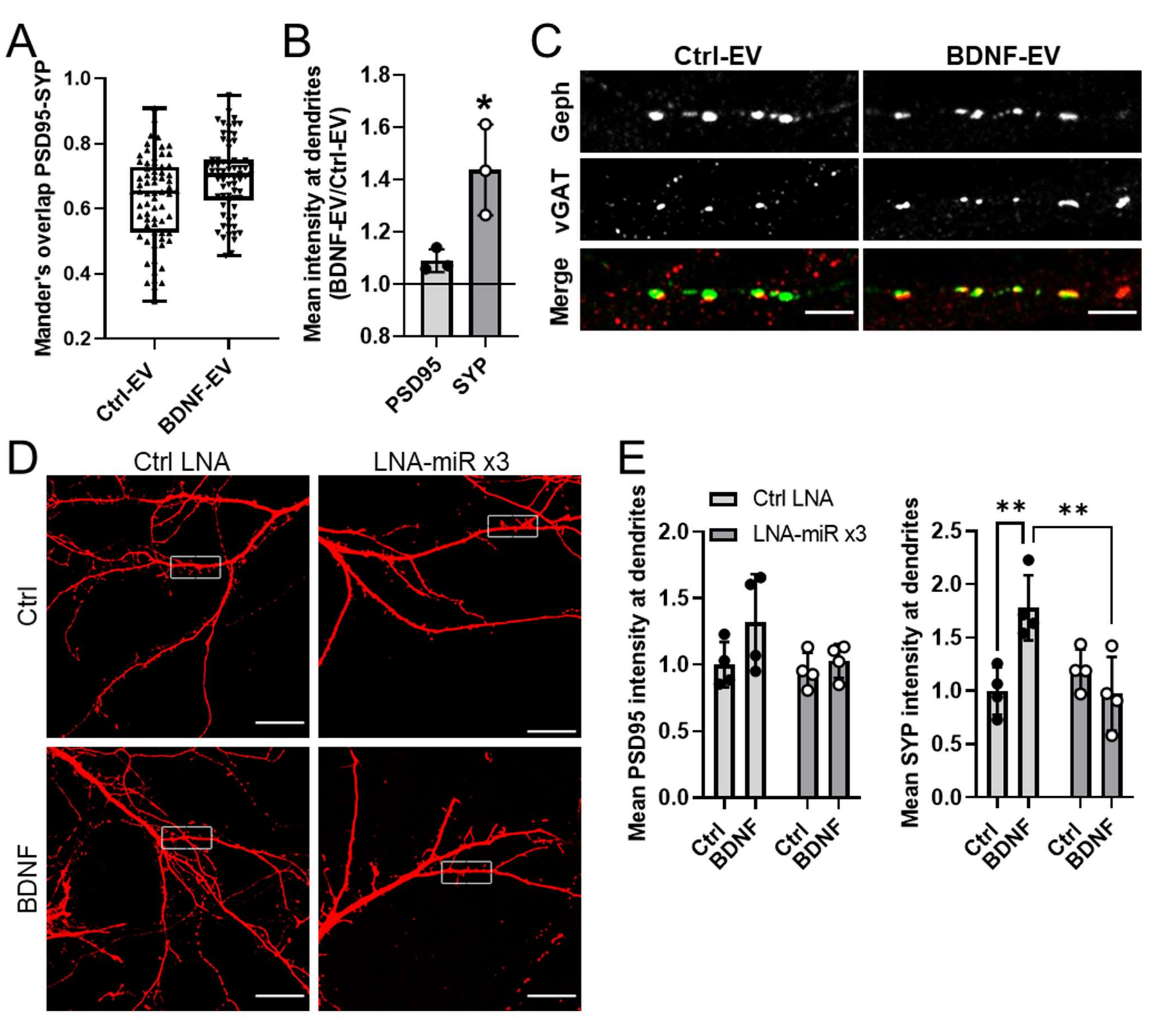


**Figure S6: BDNF-EVs regulate synapse maturation via miRNAs**

**A)** Distribution of individual mander's overlap co-efficients in neurons treated with Ctrl- or BDNF-EVs (related to **Fig. 6B**).

**B)** BDNF-EVs increase SYP intensity at neuronal dendrites. Mean PSD95 and SYP intensity was quantified in 20µm segments and values were normalized to Ctrl-EV treatment; n=3, Welch's t-test, *$p<0.05$ (related to **Fig. 6A**).

**C)** BDNF-EVs decrease the levels of Gephyrin at neuronal dendrites. Ctrl-EV or BDNF-EV treated neurons were fixed and immunostained with vGat and Gephyrin antibodies; scale bar is 5µm (related to **Fig. 6E**).

**D)** Representative images of dsRed- and LNA-expressing hippocampal neurons. Boxed insets depict selected regions of interest as shown in **figure 7A,** scale bar 20µm.

**E)** BDNF-regulated EV-miRNAs mediate the BDNF-dependent increase in SYP clustering at dendrites. The intensity of PSD95 (*left*) and SYP (*right*) in 20µm long dendrites was quantified and normalized to control, n=4, 2way ANOVA, ***p*<0.01 (related to **Fig. 7A**).

### Supplementary Data

**Table S1: List of the top 20 miRNAs regulated by BDNF in decreasing order of significance.** Shown the base mean miRNA expression and p-value with or without adjustment for multiple comparisons (Adj.)

| miR | baseMean | Fold Change | p-value | Adj. p-value |
|---|---|---|---|---|
| **mmu-miR-690 ±** | 54.65623941 | 2.759280648 | 7.70E-06 | 0.004782548 |
| **mmu-miR-132-5p ±** | 113.6137416 | 1.534084139 | 7.80E-05 | 0.024218715 |
| **mmu-miR-218-5p ±** | 6789.758564 | 1.468204261 | 0.000239291 | 0.049533179 |
| **mmu-miR-129-2-3p ±** | 698.6885011 | 0.685583457 | 0.000403781 | 0.055158418 |
| **mmu-miR-485-3p ±** | 297.4397184 | 0.770367521 | 0.00044411 | 0.055158418 |
| **mmu-miR-129-1-3p** | 106.1079428 | 0.675966927 | 0.000642137 | 0.066461185 |
| **mmu-miR-351-5p ±** | 172.4120443 | 0.667455871 | 0.001087022 | 0.096434404 |
| **mmu-miR-434-3p** | 1336.128708 | 0.808176763 | 0.001561365 | 0.121200938 |
| **mmu-miR-125a-5p** | 3281.897121 | 0.728305776 | 0.002707624 | 0.152857684 |
| **mmu-miR-6240** | 96.44859264 | 1.842111603 | 0.002621067 | 0.152857684 |
| **mmu-miR-99b-5p** | 3838.006085 | 0.793849513 | 0.002677432 | 0.152857684 |
| **mmu-miR-3068-3p** | 130.1120316 | 1.577026116 | 0.003123175 | 0.161624331 |
| **mmu-miR-329-3p** | 121.9507059 | 0.697141436 | 0.004000542 | 0.191102805 |
| **mmu-miR-132-3p ±, *N.S*** | 639.4929193 | 1.266595388 | 0.004499291 | 0.199575672 |
| **mmu-miR-92b-3p** | 633.1746534 | 0.737470926 | 0.004987532 | 0.206483833 |
| **mmu-miR-219a-2-3p** | 173.6587533 | 0.697152372 | 0.007272489 | 0.282263479 |
| **mmu-miR-151-5p** | 82.97836574 | 0.676415267 | 0.008677313 | 0.299367297 |
| **mmu-miR-296-5p** | 197.0238512 | 0.658509115 | 0.008610128 | 0.299367297 |
| **mmu-miR-1843b-5p** | 564.5156722 | 1.241956667 | 0.009357056 | 0.305828003 |

± Validated by by qPCR, N.S; not significant
