## Supplementary Methods for "Neuronal extracellular vesicles mediate BDNF-dependent dendritogenesis and synapse maturation via microRNAs"

#### List of Materials

##### Table S1: List of oligonucleotides

| Primer Name | Forward Primer | Reverse Primer |
|---|---|---|
| Mature Arc | CCCCCAGCAGTGATTCATAC | GTGATGCCCTTTCCAGACAT |
| GAPDH | AATGTGTCCGTCGTGGATCTG | CAACCTGGTCCTCAGTGTAGC |
| miR-218 binding site cloning | GGCCGCACATGGTTAGATCAAGCA; CAAGGCGCGCCACATGGTTAGATCAAGCACAAT | CATGTGGCGCGCCTTGTGCTTGATCTAACCATGTGC; CTAGATTGTGCTTGATCTAAC |
| Mycoplasma | CGCCTGAGTAGTACGTTCGC; CGCCTGAGTAGTACGTACGC; TGCCTGAGTAGTACATTCGC; TGCCTGGGTAGTACATTCGC; CGCCTGGGTAGTACATTCGC; CGCCTGAGTAGTATGCTCGC | GCGGTGTGTACAAGACCCGA; GCGGTGTGTACAAAACCCGA; GCGGTGTGTACAAACCCCGA |

| miRNA probe | Assay ID | Assay | Catalogue # |
|---|---|---|---|
| mmu-miR-132-3p | mmu480919_mir | TaqMan Advanced miRNA | A25576 |
| mmu-miR-132-5p | mmu481539_mir | TaqMan Advanced miRNA | A25576 |
| mmu-miR-218-5p | mmu481001_mir | TaqMan Advanced miRNA | A25576 |
| mmu-miR-690 | mmu482837_mir | TaqMan Advanced miRNA | A25576 |
| mmu-miR-181a-5p | mmu481485_mir | TaqMan Advanced miRNA | A25576 |
| mmu-miR-129-2-3p | mmu478544_mir | TaqMan Advanced miRNA | A25576 |
| mmu-miR-485-3p | mmu481854_mir | TaqMan Advanced miRNA | A25576 |
| mmu-miR-351-5p | mmu481806_mir | TaqMan Advanced miRNA | A25576 |
| mmu-miR-218-5p | MC10328 | miRVana miRNA mimic | 4464066 |
| mmu-miR-132-5p | MH19230 | miRVana miRNA inhibitor | 4464084 |
| mmu-miR-218-5p | MH10328 | miRVana miRNA inhibitor | 4464084 |
| mmu-miR-690 | MH11517 | miRVana miRNA inhibitor | 4464084 |
| Negative Control #1 | - | miRVana miRNA mimic | 4464058 |
| Negative Control #1 | - | miRVana miRNA inhibitor | 4464076 |

**Table S2: List of antibodies**

| Primary | Source | Application | Dilution |
|---|---|---|---|
| Actin | Sigma-Aldrich; A4700 | WB | 1:1,000 |
| AKT | Cell Signaling; #4685 | WB | 1:1,000 |
| Alix | GeneTex; GTX42812 | WB | 1:800 |
| BDNF | Abcam; ab108319 | WB | 1:1,000 |
| Calnexin | Abcam; ab2301 | WB | 1:5,000 |
| ERK1/2 (p44/42 MAPK) | Cell Signaling; #9102 | WB | 1: 1,000 |
| Flotilin-2 | Santa Cruz; sc-25507 | WB | 1: 1,000 |
| Gephyrin | Cell Signaling; #43928 | ICC | 1:800 |
| GFAP | Synaptic Systems; #173004 | ICC | 1:500 |
| Grp75 | Abcam; ab53098 | WB | 1:1,000 |
| Lamp1 | Abcam; ab24170 | WB | 1:1,000 |
| | | ICC | 1:100 |
| MAP2 | Abcam; ab11268 | ICC | 1:1,000 |
| N-Cadherin | BD Biosciences; #610920 | WB | 1:1,000 |
| Phospho-AKT (Ser473) | Cell Signaling; #9271 | WB | 1:2,000 |
| Phospho-ERK1/2 (Thr202/Tyr204) | Cell Signaling; #q4370 | WB | 1:1,000 |
| Phospho-TrkB (Tyr816) | Merckmillipore; ABN1381 | WB | 1:500 |
| PSD95 | Merckmillipore; MAB1596 | ICC | 1:100 |
| Synaptophysin | Abcam; ab16659 | ICC | 1:100 |
| TrkB | RnD Systems; AF1494 | WB | 1:500 |
| TSG101 | Santa Cruz; sc-7964 | WB | 1:500 |
| vGAT | ThermoScientific; MA5-24643 | ICC | 1:100 |
| **Secondary** | **Source** | **Application** | **Dilution** |
| Alexa-Fluor conjugated | Invitrogen | ICC | 1:2,000 |

| | | | |
|---|---|---|---|
| HRP-conjugated | JacksonImmuno | WB | 1:4,000 |

*IB; immunoblotting, ICC; immunocytochemistry*

### Supplementary Methods

#### EV isolation

*Differential ultracentrifugation (UC)*

Unless otherwise stated, all steps were carried out on ice or at 4˚C. Collected cell culture supernatants were centrifuged as depicted in **Fig. 1A**. For mEV isolation, 10,000 x*g* pellets were washed once in cold PBS and centrifuged again for 30min at 10,000x*g*. For sEV isolation, 10,000x*g* supernatants were transferred to thickwall polyallomer tubes (Beckman Coultier) and subjected to UC for 1h at 100,000x*g* in a fixed-angle rotor (rotor; TLA 100.3, *k*-factor; 48, Optima MAX-XP tabletop ultracentrifuge, Beckman Coulter). sEV pellets were washed in cold PBS by vortexing before the second UC step. UC supernatants were discarded by decanting. EV pellets were re-suspended in 20mM HEPES-buffered saline (pH7.4) containing 0.025% Tween-20 (Hepes-t) and protease inhibitors by gentle shaking for 20 min at room temperature

For fluorescent labeling of EVs, 100,000 x*g* pellets from the first UC step were re-suspended in HEPES-t and incubated with 100nM lipilight (Idylle) for 3 min, after which EVs were diluted 15-fold in PBS and pelleted again by UC. No-cell controls were processed in parallel, using non-conditioned cell culture medium.

*Size Exclusion Chromatography (SEC)*

Isolation of sEVs by SEC was performed using 10,000 x*g* supernatants (**Fig. 1A**) and qEVoriginal (70nm) columns (IZON Bioscience), according to manufacturer's instructions. Fractions of interest were pooled together and concentrated using 3 kDa ultra-filtration devices (Amicon).

#### Nanoparticle tracking analysis (NTA)

NTA was performed on NanoSight NS500 LM10 instrument and a LM14 viewing unit equipped with a 532nm laser (NanoSight Ltd). Five 60-second measurements per sample were used for analysis using NTA 2.3 software. Values were normalized to relative protein concentration of donor neurons.

#### Scanning transmission electron microscopy (STEM) and analysis

Glow-discharged formvar/carbon-coated TEM grids (400 mesh, copper) were floated on 20µL drops of sEV samples for 10 minutes. After removing excess solution, the grids were floated on 2% uranyl acetate for 30 min. Afterwards the staining solution was removed by dabbing the grids onto filter paper. TEM grids were further dried for at least 1 hour before imaging on a Gemini II SEM column (Zeiss Crossbeam 550) at 30kV and 150pA (high resolution imaging mode) with a STEM detector. Quantification of particle size and number was performed in ImageJ particle analyzer using thresholded images. Parameters were set to exclude particles less than 75nm or present at the edges of the field of view.

#### Protein analysis

Donor cells were washed thrice in cold PBS and lysed in CHAPS buffer (1% CHAPS, 5mM EDTA and 50mM Tris-HCl (pH8)). Post-nuclear supernatants were isolated by centrifugation at 4,500x*g* for 10min at 4˚C. For quantification of protein concentration, EV pellets were lysed in CHAPS buffer (EVs from 100,000 donor cells / µl CHAPS) by vortexing at full speed for 2 min and cell and EV lysates were subjected to Bicinchonic acid (BCA) assay.

#### Immunoblotting

Cell lysates or EV pellets were re-suspended in Laemmli sample buffer by vortexing, and heated at 70˚C for 10 min before being subjected to SDS-PAGE electrophoresis. Proteins were transferred onto polyvinylidine difluoride (PVDF) membranes, which were blocked in Tris-buffered saline containing 0.1% Tween-20 and either 4% semi-skimmed milk powder or 5% BSA for phospho-specific antibodies. Primary and secondary antibodies were diluted in blocking buffer (see also Table S2). Blots were detected using the ChemiDoc MP imaging system (Bio-rad).
